# Autosomal recessive loci contribute significantly to quantitative variation of male fertility in a dairy cattle population

**DOI:** 10.1101/2020.12.11.421354

**Authors:** Maya Hiltpold, Naveen Kumar Kadri, Fredi Janett, Ulrich Witschi, Fritz Schmitz-Hsu, Hubert Pausch

**Author notes:** **Corresponding author** Correspondence to Maya Hiltpold.

## Abstract

Cattle are ideally suited to investigate the genetics of male fertility. Semen from individual bulls is used for thousands of artificial inseminations for which the fertilization success is monitored. In a cohort of 3881 bulls that had genotypes at 589,791 SNPs, we reveal four novel recessive QTL for male fertility using haplotype-based association testing. We detect either missense or nonsense variants in *SPATA16*, *VWA3A, ENSBTAG00000006717* and *ENSBTAG00000019919* that are in linkage disequilibrium with the QTL. A QTL for bull fertility on BTA1 is also associated with sperm head shape anomalies. Using whole-genome sequence and transcriptome data, we prioritise a missense variant (p.Ile193Met) in *SPATA16* as candidate causal variant underlying this QTL. Our findings in a dairy cattle population provide evidence that recessive variants may contribute substantially to quantitative variation in male fertility in mammals.

## Introduction

Male fertility is a complex trait that is determined by genetic and non-genetic sources of variation. Because of its low heritability, a large sample size is required to investigate the genetic architecture of male fertility. Large cohorts of males with repeated measurements for reproductive traits are not available in many species including humans, but are accessible in cattle where semen samples from individual bulls are used for thousands of artificial inseminations ^1^. Because these data facilitate the disentanglement of male and female factors contributing to establishing pregnancy ^2^, the fertility of bulls can be quantified objectively.

Semen quality has a large effect on insemination success ^3–5^. On entering the semen collection center, the semen quality of each bull is examined as part of an in-depth breeding soundness evaluation ^6^. The semen quality of artificial insemination bulls varies over time, consequently the fertilization rates may differ substantially between ejaculates. In order to ensure high and uniform insemination success, the volume, sperm concentration, sperm head and tail morphology, and sperm motility is examined in all ejaculates immediately after semen collection. Ejaculates that fulfill predefined quality requirements are diluted and filled into doses that contain between 15 and 25 million sperm, depending on semen quality.

Phenotypes describing male reproductive traits and dense microarray-derived genotypes for many artificial insemination sires enable genome-wide association testing. Quantitative trait loci (QTL) for bull fertility have been detected in Holstein ^7–10^ and Jersey ^11^ cattle. Nevertheless, these studies did not attempt to reveal the causal variants underlying the QTL due to the use of sparse marker panels. Moreover, the sample sizes were relatively small providing low statistical power to detect QTL for a trait with low to moderate heritability like bull fertility. Recent investigations provide growing evidence that non-additive effects contribute to inherited variation in bull fertility ^8,12–15^. However, the underlying genetic mechanisms remain largely unknown.

Here, we apply haplotype-based association testing to detect QTL for bull fertility and semen quality in a large mapping cohort of bulls from the Brown Swiss cattle breed that had dense microarray-derived genotypes. Our association studies revealed five recessive QTL for bull fertility. Using two complementary sequence-based fine-mapping approaches and transcriptome data, we identify candidate causal variants in coding sequences of genes that are expressed in the male reproductive tract.

## Results

To detect QTL for bull fertility, we considered 3736 Brown Swiss (BSW) bulls, that were kept at semen collection centers in Switzerland, Germany and Austria. Bull fertility was quantified based on artificial insemination success adjusted for confounding factors (see Methods). The bulls had partially imputed genotypes at 589,791 autosomal SNPs with minor allele frequency greater than 0.5 %. Genomic restricted maximum likelihood estimation indicated that the autosomal markers explained 0.10 ± 0.02% of the phenotypic variance in bull fertility.

### Recessive QTL for bull fertility are located on chromosomes 1, 6, 18, 25 and 26

Haplotype-based association tests that were based on an additive mode of inheritance revealed three QTL located on chromosomes 1, 6, and 26 for bull fertility that exceeded the Bonferroni-corrected genome-wide significance threshold of 2.68 × 10^−7^ (Table 1, Figure 1a). We previously showed that the QTL for bull fertility at BTA6 is in linkage disequilibrium with the recessive BTA6:58,373,887 T-allele (rs474302732) that activates cryptic splicing in *WDR19,* resulting in reduced semen quality ^13^. In order to investigate the association of the BTA6:58,373,887 T-allele in our mapping cohort, we imputed the genotypes of the 3736 bulls to the whole-genome sequence level using a breed-specific reference panel of 368 sequenced cattle. A sequence-based association analysis that was based on a recessive mode of inheritance confirmed that rs474302732 is strongly associated (P = 2.72 × 10^−45^) with bull fertility. The association with bull fertility was stronger for rs474302732 than the most significantly associated haplotype (Table 1). Four non-coding variants (57,900,948 bp, 57,950,075 bp, 57,408,389 bp, 57,810,181 bp) had slightly lower P values (P = 5.36 × 10^−46^ - 3.81 × 10^−47^) than rs474302732. Due to its large effect on male fertility, we subsequently fixed the top haplotype of the BTA6 QTL as a covariate in the association model. When the top haplotype of the BTA6 QTL was fixed as a covariate in the model, the association signal at BTA6 disappeared (Figure 1b). In the conditional analysis, the P value of the top-associated haplotype at the BTA1 QTL was slightly above the Bonferroni-corrected significance threshold (Figure 1b). The QTL on BTA26 remained significant (P = 3.25 × 10^−9^).

**Table 1:**
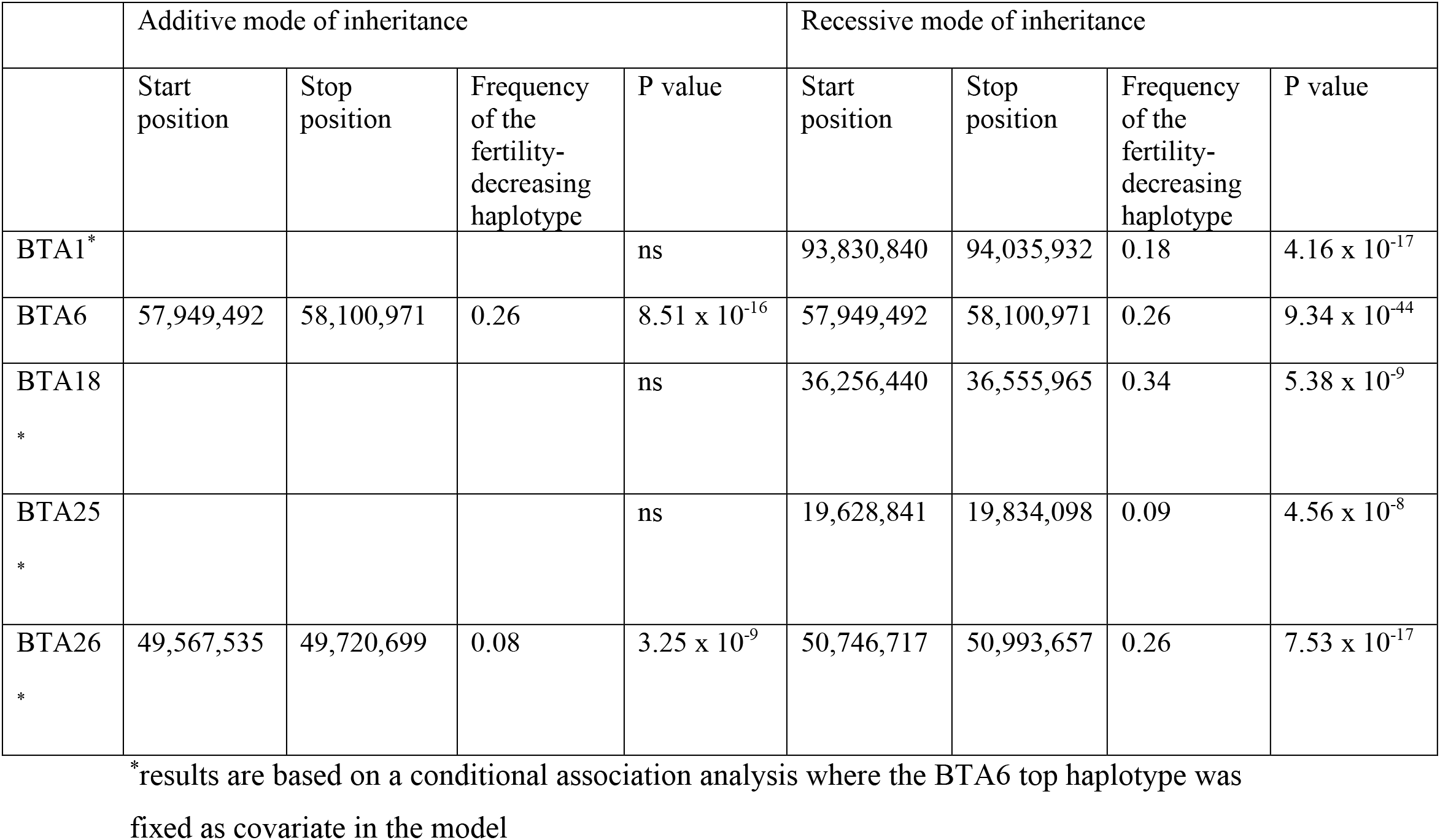
Position of five recessive QTL for male fertility in BSW cattle.

|  | Additive mode of inheritance |  |  |  | Recessive mode of inheritance |  |  |  |
| --- | --- | --- | --- | --- | --- | --- | --- | --- |
|  | Start position | Stop position | Frequency of the fertility-decreasing haplotype | P value | Start position | Stop position | Frequency of the fertility-decreasing haplotype | P value |
| BTA1* | | | | ns | 93,830,840 | 94,035,932 | 0.18 | $4.16 \times 10^{-17}$ |
| BTA6 | 57,949,492 | 58,100,971 | 0.26 | $8.51 \times 10^{-16}$ | 57,949,492 | 58,100,971 | 0.26 | $9.34 \times 10^{-44}$ |
| BTA18* | | | | ns | 36,256,440 | 36,555,965 | 0.34 | $5.38 \times 10^{-9}$ |
| BTA25* | | | | ns | 19,628,841 | 19,834,098 | 0.09 | $4.56 \times 10^{-8}$ |
| BTA26* | 49,567,535 | 49,720,699 | 0.08 | $3.25 \times 10^{-9}$ | 50,746,717 | 50,993,657 | 0.26 | $7.53 \times 10^{-17}$ |

\*results are based on a conditional association analysis where the BTA6 top haplotype was fixed as covariate in the model

**Figure 1:**
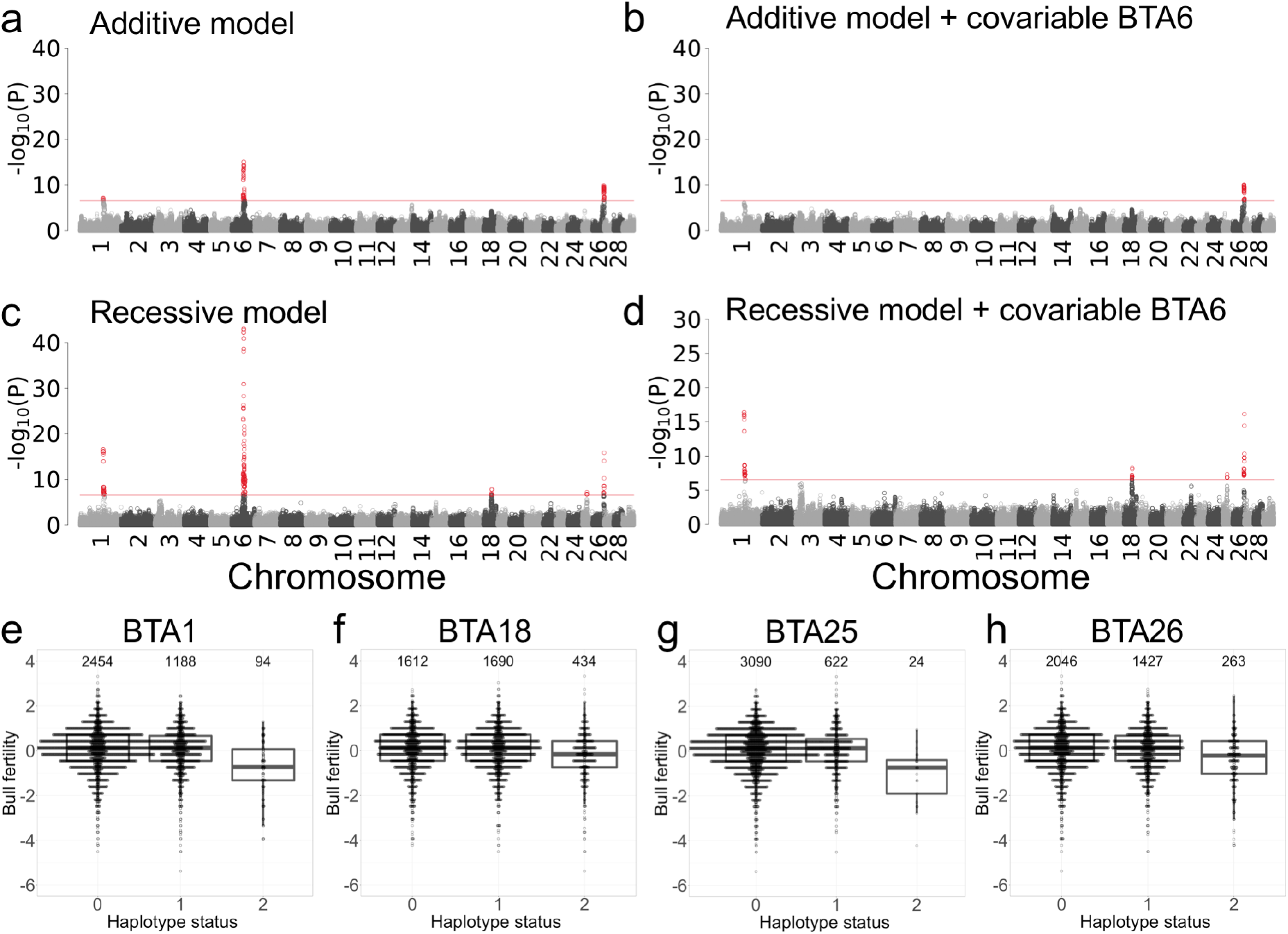
Results of haplotype-based genome-wide association studies with bull fertility. Manhattan plots representing the association (−log_10_(P)) of 186,278 haplotypes with bull fertility in 3736 Brown Swiss bulls (**a-d**). Association tests were performed based on either additive (**a, b**) or recessive (**c, d**) modes of inheritance. Results are presented before (**a, c**) and after (**b, d**) accounting for a QTL at BTA6. Red dots represent haplotypes that exceed the Bonferroni-corrected significance threshold (red horizontal line). Effect of the top haplotypes at BTA1 (**e**), BTA18 (**f**), BTA25 (**g**) and BTA26 (**h**) on bull fertility (0-non-carrier, 1-heterozygous, 2-homozygous). The values above the boxplots indicate the number of bulls carrying 0, 1 and 2 copies of the top haplotype.

Next, we repeated the haplotype-based association test assuming a recessive mode of inheritance. The association with bull fertility was more pronounced (i.e., the P values of the top haplotypes were lower) at the QTL on chromosomes 1, 6 and 26 (Table 1, Figure 1c & d). Moreover, the recessive model revealed two QTL at chromosomes 18 and 25 that were not detected using the additive model. Visual inspection of the haplotype effects corroborated that the four novel QTL detected on chromosomes 1, 18, 25, and 26 act in a recessive manner (Figure 1e, f, g, h). Compared to non-carrier and heterozygous haplotype carrier bulls, the fertility of bulls that carry the BTA1, BTA18, BTA25 and BTA26 top haplotype in the homozygous state is reduced by 0.80 ± 0.10, 0.28 ± 0.05, 1.09 ± 0.20 and 0.52 ± 0.06 phenotypic standard deviations, respectively. A linear regression analysis conditional on the top 10 principal components and the top haplotype at the BTA6 QTL confirmed that fertility does not differ between heterozygous and non-carrier bulls at the four novel QTL (P_BTA1_ = 0.29, P_BTA18_ = 0.61, P_BTA25_ = 0.79, P_BTA26_ = 0.59), confirming their recessive inheritance.

Between 24 and 434 bulls were homozygous carriers of fertility-decreasing haplotypes at the four novel and one previously detected QTL. Across the five QTL, 2827, 604, 259, 44, and 2 bulls were homozygous carriers of 0, 1, 2, 3, and 4 top haplotypes, respectively. In order to quantify the impact of the five recessive QTL on insemination success in the BSW population, we investigated the 56-day non-return rate (NRR56; mean = 65.06 ± 4.53 %) in cows inseminated with semen from 1322 BSW bulls, estimated from at least 300 inseminations per bull. The NRR56 is the proportion of cows that is not re-inseminated within a 56-day interval after the first insemination and is considered as a proxy for the insemination success.. The mean NRR56 is 0.5 phenotypic standard deviations higher (P = 2.75 × 10^−8^, two-tailed t-test) in 1000 bulls that do not carry any of the top haplotypes in the homozygous state than in 322 bulls that are homozygous for at least one of the five recessive QTL (65.48 ± 4.33 vs. 63.76 ± 4.89 %).

### A bull fertility QTL on BTA1 affects sperm head morphology

A recessive QTL for bull fertility is located on BTA1. The top haplotype (P = 4.16 x 10^−17^) resides between 93,830,840 and 94,035,932 bp. The fertility-decreasing haplotype occurs at a frequency of 0.18 in the BSW population. We detected 94 and 1188 bulls that carried the top haplotype in the homozygous and heterozygous state, respectively.

In order to unravel the mechanism through which the QTL impacts male fertility, we analysed routinely collected semen quality data that were available for 32 homozygous, 372 heterozygous and 498 non-carrier bulls. The BTA1 QTL was not associated with ejaculate volume (P = 0.07), sperm concentration (P = 0.19), sperm motility (P = 0.10), sperm head morphology (P = 0.53) or sperm tail morphology (P = 0.19, Supplementary Figure 1) assessed in fresh semen. However, the insemination straws contained more (+1.57 million, P = 4.78 × 10^−6^, Supplementary Figure 1f) sperm in homozygous than heterozygous and non-carrier bulls, which indicates an attempt to compensate for low sperm quality by providing additional sperm per straw, although the association with the number of sperm per straw was above the Bonferroni-corrected significance threshold (P_Bonf_ = 2.68 × 10^−7^). Moreover, more ejaculates were discarded due to insufficient semen quality from homozygous than heterozygous and non-carrier bulls (14.8 vs. 6.5 %).

The association pattern of the BTA1 top haplotype was puzzling. Homozygosity for the top haplotype compromises bull fertility but not routinely recorded fresh semen quality. However, the insemination doses of homozygous bulls contain more sperm per straw. In order to further examine this apparent contradiction, we studied 1577 spermiograms that were collected as part of the andrological examination of 575 prospective artificial insemination bulls. During the andrological examination, sperm morphology of at least one ejaculate per bull is systematically evaluated, before the semen is used for artificial inseminations. This evaluation is repeated either until the semen quality meets predefined quality thresholds or the bull is rejected for artificial insemination due to insufficient semen quality. Spermiogram features were provided for between 1 and 13 (median = 2) ejaculates of 575 Swiss BSW bulls. From the spermiogram reports, we derived 21 sperm quality phenotypes (see Methods)

Genome-wide haplotype-based association testing revealed that the QTL at BTA1 affects four (out of 21) sperm morphology features. Bulls that are homozygous for the top haplotype produce ejaculates that contain less normal spermatozoa (−10.6 %, P = 1.72 × 10^−10^), twice as many sperm with non-compensatory defects (+5.4 %, P = 9.07 × 10^−16^), 1.5 times more sperm with major defects (+8.9 %, P = 2.55 × 10^−11^) and twice as many sperm with head shape anomalies (+5.6 %, P = 3.98 × 10^−17^) (Figure 2, Supplementary Figure 1h, i, l, & m). These data suggest that an increased proportion of sperm with abnormal head morphology compromises bull fertility, as no other sperm morphology feature was affected by the BTA1 QTL. The number of sperm morphology evaluations was also higher (P = 1.63 × 10^−10^) for homozygous than heterozygous and non-carrier bulls (Supplementary Figure 1g). Eventually, only 21 out of 34 homozygous bulls (61.7 %) compared to 520 out of 561 (92.7 %) heterozygous and non-carrier bulls passed the sperm morphology examination and produced ejaculates that are suitable for breeding (≥ 65 % normal sperm and ≤ 20 % non-compensatory defects). Sperm morphology did not differ between heterozygous haplotype carriers and non-carrier bulls, corroborating recessive inheritance.

**Figure 2:**
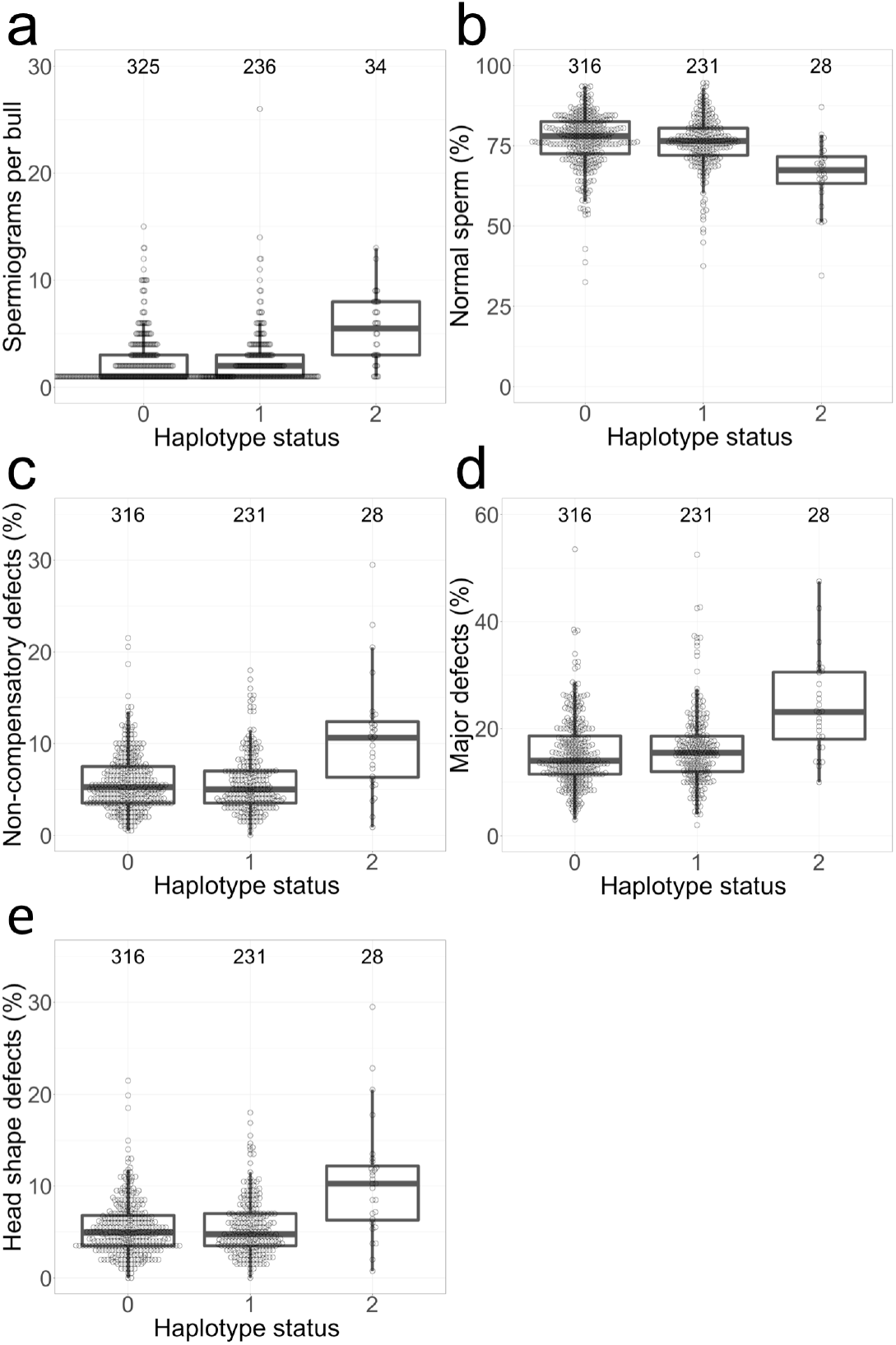
Effect of the BTA1 top haplotype on sperm morphology. Boxplots representing the sperm morphology of non-carrier, heterozygous, and homozygous (haplotype status 0, 1, and 2) bulls. The numbers above the boxplots indicate the number of bulls in the respective group. Compared to non-carrier and heterozygous bulls, **a** the number of sperm morphology examinations is twice as high (5.32 ± 3.25 vs. 2.49 ± 2.50), **b** the amount of normal spermatozoa is ~10 percentage points lower (66.19 ± 10.43 % vs. 76.45 ± 8.51 %), **c** the amount of sperm with non-compensatory defects is ~ 5 percentage points higher (10.61 ± 6.28 % vs. 5.79 ± 3.19 %), **d** the amount of sperm with major defects is ~ 8 percentage points higher (24.08 ± 8.99 % vs. 15.70 ± 6.82 %), **e** and the amount of spermatozoa with head shape anomalies is ~ 5 percentage points higher (10.49 ± 6.30 % vs. 5.42 ± 3.09 %) in homozygous bulls.

We examined and imaged fresh sperm from two bulls homozygous for the top haplotype using oil-immersion phase-contrast light microscopy. At semen collection, the bulls were 644 and 504 days old. Some spermatozoa had morphological anomalies (i.e., rounder and shorter heads, heads with rounder frontal part (pear-shape /pyriform), narrower head base (tapered), and heads with abnormal contour (uneven shaped)) that are classified as head shape anomalies in the spermiogram examination (Supplementary Figures 2 & 3). However, the vast majority of the spermatozoa (85 and 84.6 %) were normal. Acrosome defects were not apparent.

### A missense variant in and non-coding variants upstream *SPATA16* are associated with the BTA1 QTL

In order to detect candidate causal variants for the increased number of abnormal sperm and reduced fertility associated with the BTA1 QTL, we applied a two-step fine-mapping approach. First, we used whole-genome sequence data of 125 BSW bulls to identify variants that are compatible with the inheritance of the top haplotype. The average fold coverage of the sequenced bulls was 10.1 ± 4.3-fold. The BTA1 QTL top haplotype was determined for the sequenced bulls based on their microarray-derived genotypes. Second, we imputed whole-genome sequence variant genotypes for the 3736 BSW bulls of the mapping cohort using 368 reference animals to perform an association study between imputed sequence variant genotypes and bull fertility.

One and 40 sequenced bulls carried the top haplotype in the homozygous and heterozygous state, respectively. Of 44,948 sequence variants that were polymorphic within a window encompassing 3 Mb on either side of the top haplotype, we detected 764 variants between 93,614,265 and 96,742,540 bp that were compatible with recessive inheritance, i.e. these variants were heterozygous in haplotype carriers and homozygous in the bull that carried the haplotype in the homozygous state. The sequence coverage did not differ between heterozygous, homozygous and non-carrier bulls within that interval.

Of the 764 compatible variants, 505 were also significantly associated with bull fertility at the genome-wide Bonferroni-corrected significance threshold of 3.89 × 10^−9^ (Figure 3a). The significantly associated variants clustered between *SPATA16* encoding spermatogenesis associated protein 16 and *NLGN1* encoding neuroligin (Figure 3a). The most significantly associated variant (BTA1:93,972,058G>A, rs379712951) is in an intron of *NLGN1*, 373 kb upstream the translation start site of *SPATA16*, within the top window from the haplotype-based association study. The P value is slightly lower for the BTA1:93,972,058G>A variant than for the most significantly associated haplotype (P=3.00 × 10^−17^ vs. 4.16 × 10^−17^). We did not consider *NLGN1* as a candidate gene for an impaired bull fertility, as it is not notably expressed in adult bull testis (TPM < 1). However, we considered *SPATA16* as a positional and functional candidate gene for the BTA1 QTL, because it is testes-specific expressed in cattle (http://cattlegeneatlas.roslin.ed.ac.uk) and human (https://gtexportal.org/home/gene/SPATA16). Moreover, *SPATA16* mRNA is highly abundant in testis tissue of adult bulls (TPM = 295) (Figure 3b).

**Figure 3:**
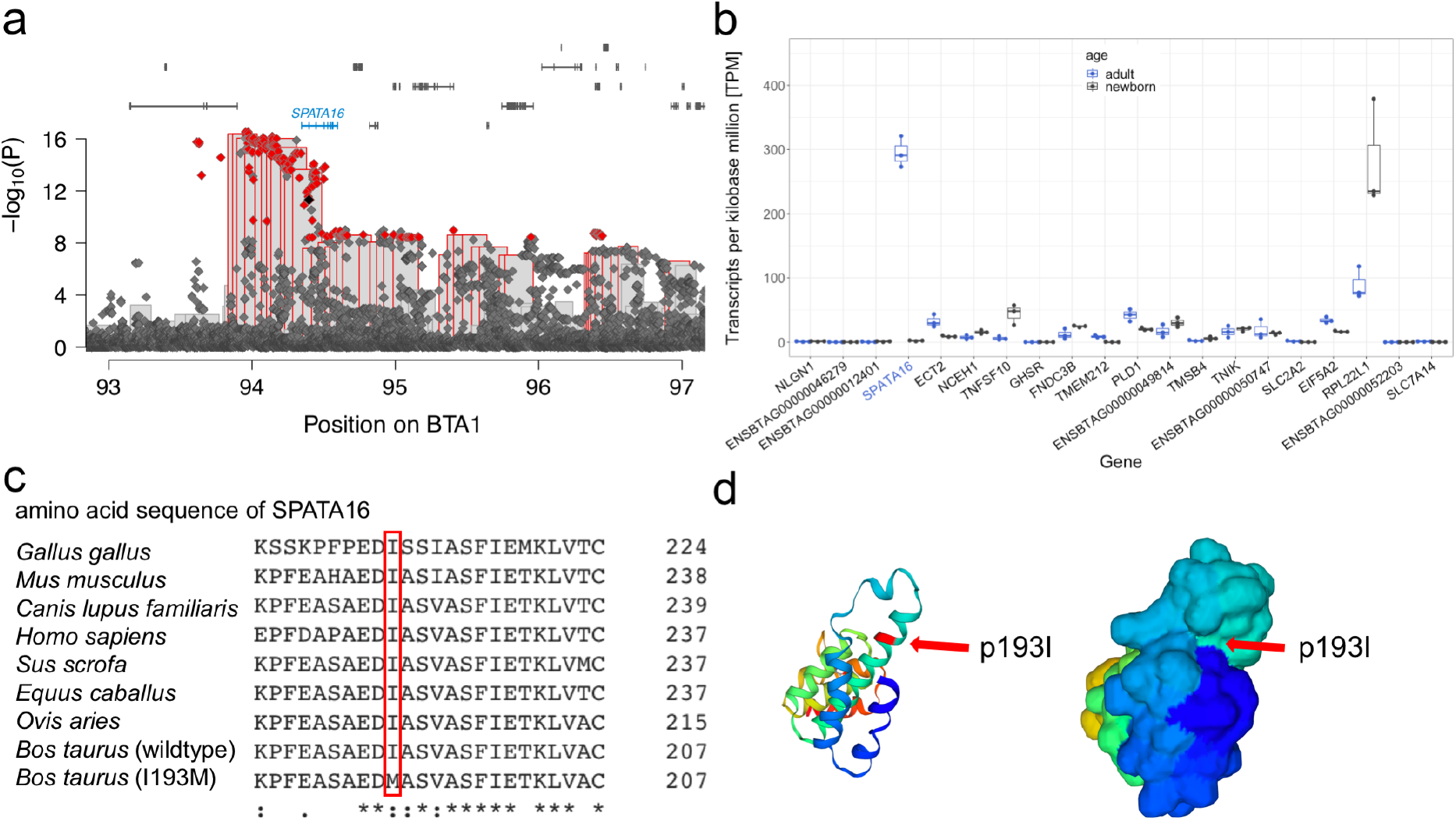
Fine mapping of a QTL for male fertility on BTA1. **a** Association of haplotypes (bars) and imputed sequence variants (diamonds) located between 93 and 97 Mb with bull fertility. Red framed bars represent significantly associated haplotypes. Imputed sequence variants that exceeded the Bonferroni-corrected significance threshold and were compatible with recessive inheritance of the top haplotype are displayed in red. The black dot indicates the SPATA16:p.Ile193Met variant (BTA1:94,396,804A>G). Blue colour indicates testis-specific expressed genes. **b** Transcript abundance (quantified in transcripts per million (TPM)) in testis tissue of adult (blue) and newborn cattle. Blue labels indicate testis-specific expressed genes. **c** Clustal Omega multi-species alignment of SPATA16 in *Gallus gallus* (ENSGALT00000052301.3)*, Mus musculus* (ENSMUST00000047005.10)*, Canis lupus familiaris* (ENSCAFT00000062886.1), *Homo sapiens* (ENST00000351008.4), *Sus scrofa* (ENSSSCT00000060926.2), *Equus caballus* (ENSECAT00000012917.2), *Ovis aries* (ENSOART00020000622.1) and *Bos taurus* wild-type (ENSBTAT00000061217.3) and mutant *(I193M)*. **d** TrEMBL 3D-structure prediction of wildtype bovine SPATA16 (F1MN96) in cartoon (left) and surface (right) representation. The isoleucine at position 193 (red arrow) resides within an alpha helix on the surface of SPATA16.

Two of the significantly associated variants compatible with recessive inheritance were in coding regions: a missense variant in *SPATA16* (rs440830663 at BTA1:94,396,804A>G, ENSBTAP00000053460.3:p.Ile193Met, P = 4.83 × 10^−12^) and a synonymous variant in *GHSR* encoding growth hormone secretagogue receptor (rs714884352, BTA1:95,029,804C>T, ENSBTAP00000014446.5:p.228Leu, P = 3.7 x 10^−9^). Pathogenic alleles of human *SPATA16* are associated with globozoospermia, i.e., round-headed, often acrosome lacking spermatozoa (Dam et al., 2007; ElInati et al., 2016). The p.Ile193Met variant compatible with recessive inheritance resides in an evolutionarily conserved tetratricopeptide repeat domain of SPATA16 (Figure 4c). The isoleucine at position 193 is on the surface of the protein (Figure 4d). According to protein structure modelling with *Missense3D*, a methionine at position 193 does not alter the tertiary structure of SPATA16. However, it is predicted to be deleterious to SPATA16 function (SIFT score: 0.03, PolyPhen-2 score: 0.998). The P value is lower and the effect on male fertility (−0.80 vs. −0.57) is larger for the most significantly associated intergenic than the p.Ile193Met variant in *SPATA16*. When the SPATA16:p.Ile193Met variant was fixed as covariate in the haplotype and sequence-based association model (Supplementary Figure 4b), the original top haplotype and the variants upstream *SPATA16* were still associated with bull fertility, albeit not at the Bonferroni-corrected significance threshold. The P values of the most significantly associated variant (BTA1:93,972,058 bp) and of the top haplotype from the conditional analysis were 1.52 × 10^−6^ and 1.74 × 10^−6^. When the association analysis was conditioned on the BTA1:93,972,058 bp variant, i.e., the most significantly associated variant from the sequence-based association study, the QTL signal was absent in both association studies (Supplementary Figure 4c). In our set of partially imputed sequence variants, BTA1:94,396,804A>G and BTA1:93,972,058G>A were in high linkage disequilibrium (r^2^ = 0.858).

**Figure 4:**
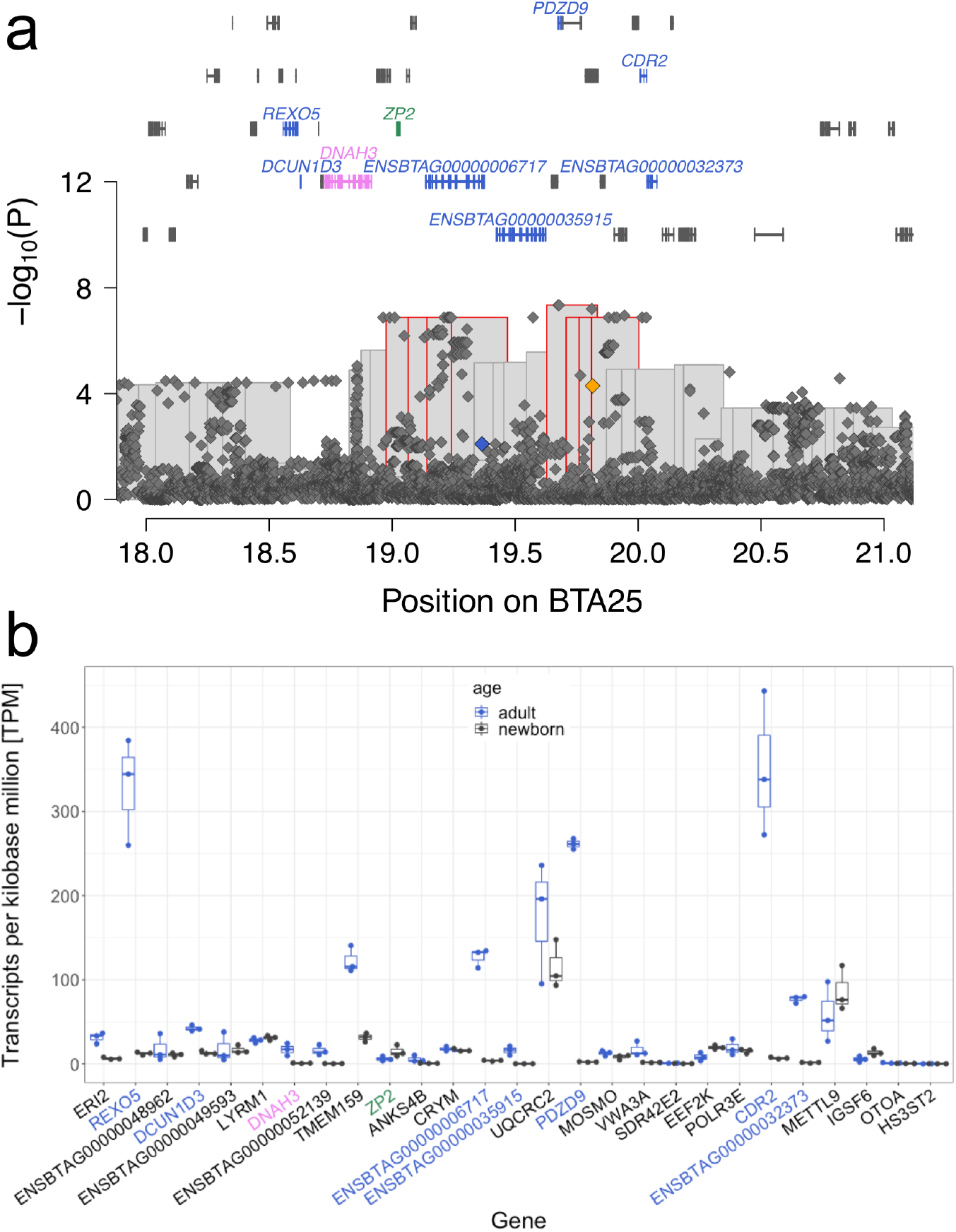
Detailed view of a QTL for bull fertility on BTA25. **a** Manhattan plot representing the association of imputed sequence variants (diamonds) and haplotypes (bars) with bull fertility. Red framed bars represent haplotypes that met the Bonferroni-corrected significance threshold. The orange and blue diamonds represent the nonsense variant (p.Arg505Ter) in *VWA3A* and frameshift variant (p.Ile1167LeufsTer) in *ENSBTAG00000006717*. Symbols over the Manhattan plot indicate genes and transcripts that are annotated at the QTL region. Genes and transcripts with testis-biased expression, male and female reproductive tract specific expression, and testis and brain specific expression, are indicated in blue, violet and green colour, respectively. **b**Transcript abundance quantified in TPM in testis tissue of adult (blue) and newborn cattle. Genes and transcripts with testis-biased expression, male and female reproductive tract specific expression, and testis and brain specific expression, respectively, are indicated in blue, violet and green colour, respectively.

### Coding variants in *ENSBTAG00000006717* and *VWA3A* segregate with the BTA25 QTL

A haplotype-based association analysis conditional on the BTA6 QTL yielded eight significantly associated haplotypes on BTA25 located between 18,975,561 and 20,002,993 bp. The top haplotype (19,628,841 – 19,834,098 bp, P=4.56 × 10^−8^) has a frequency of 9 % in the BSW population. Despite its strong effect (−1.09 ± 0.20 standard deviations) on bull fertility, the top haplotype was neither associated with routinely examined semen quality nor with sperm morphology features that were assessed as part of the andrological examination. Of 125 sequenced BSW cattle, 1 and 20 carry the top haplotype in the homozygous and heterozygous state, respectively. Sequence coverage does not differ between homozygous, heterozygous and non-carrier bulls at the BTA25 QTL. We detected 48,121 sequence variants within ±3 Mb of the top haplotype, of which 778 were compatible with the recessive inheritance of the top haplotype. Of the compatible variants, six reside in protein-coding sequences: four synonymous variants in *ZP2* (rs110876106, BTA25:19,021,021C>T, p.Pro630%3D), *VWA3A* (rs209406752, BTA25:19,788,285C>T, p.Leu33%3D; rs210416159, BTA25:19,798,888C>T, p.Asp32%3D), and *EEF2K* (rs110751593, BTA25:19,941,219C>T, p.Cys538%3D), a frameshift variant in *ENSBTAG00000006717* (BTA25:19,365,282CA>C, ENSBTAP00000054375.1:p.Ile1167LeufsTer) and a stop gained variant in *VWA3A* (rs434854120, BTA25:19,814,925C>T, ENSBTAP00000021610.1:p.Arg505Ter). Due to a putatively high impact on protein function, we considered the stop gained and frameshift variants in *VWA3A* and *ENSBTAG00000006717* as candidate causal variants for the BTA25 QTL.

*ENSBTAG00000006717* encoding the nondescript ATP-binding cassette sub-family A member 3-like protein is highly expressed in testis tissue of adult bulls (127 TPM) and shows a testis-specific expression in cattle ^16^ (http://cattlegeneatlas.roslin.ed.ac.uk). The frameshift partly truncates an evolutionarily conserved domain “ATPase associated with a variety of cellular activities (AAA)”. *VWA3A* encoding von Willebrand factor A domain-containing protein 3A is expressed in testis tissue of adult bulls at 17 TPM. The stop mutation truncates VWA3A by 58%. The von Willebrand factor A domain-containing protein 3A is a motile cilia-associated protein and might contribute to the beating movement of the flagella ^17^. An association study between imputed sequence variants and bull fertility revealed no variants exceeding the Bonferroni-corrected significance threshold at the BTA25 QTL (Figure 4a). However, 38 variants with P < 7.66 × 10^−7^ (i.e., P values that are slightly above the significance threshold) were between 18,963,390 and 20,033,799 bp. The most significantly associated variant (P = 4.56 × 10^−8^) was at 19,675,705 bp in an intron of *PDZD9* encoding PDZ domain containing 9. The P values were considerably larger for the frameshift and nonsense variant in *ENSBTAG00000006717* (P=7.98 × 10^−3^) and *VWA3A* (P=5.11 × 10^−5^), respectively.

### A missense variant in *ENSBTAG00000019919* segregates with the BTA26 QTL

The top haplotype (P = 4.9 × 10^−17^) at the BTA26 QTL is between 50,746,717 and 50,993,657 bp. The frequency of the top haplotype in the BSW population is 0.26. Our mapping cohort contained 263 homozygous and 1427 heterozygous haplotype carriers. Of 125 sequenced BSW bulls, 11 and 63 carried the top haplotype in the homozygous and heterozygous state, respectively. The top haplotype was not associated with any other semen quality or sperm morphology trait.

Twenty-six non-coding and two coding variants that were located between 50,748,638 and 50,979,057 bp were compatible with the top haplotype. A synonymous (BTA26:50,857,552C>T, rs381031945, ENSBTAP00000026536.1:p.831Asp%3D) and a missense variant (BTA26:50,850,915C>G, rs378141069, ENSBTAP00000026536.1:p.Asn576Lys) in *ENSBTAG00000019919* were compatible with the top haplotype. The Ensembl and Refseq annotations indicate that *ENSBTAG00000019919* is the bovine ortholog of *CFAP46* encoding cilia and flagella associated protein 46. The two coding variants in *ENSBTAG00000019919* were in complete linkage disequilibrium and highly significantly associated with bull fertility (P=1.3 × 10^−15^) in the sequence-based association study (Figure 5a). The p.Asn576Lys-variant is predicted to be deleterious (SIFT: 0.02, PolyPhen-2: 0.921) to protein function. *ENSBTAG00000019919* is expressed at 21 TPM in adult bull testis (Figure 5b). In *Chlamydomonas reinhardtii*, a CFAP46 homolog is essential for normal flagellar motility ^18^.

**Figure 5:**
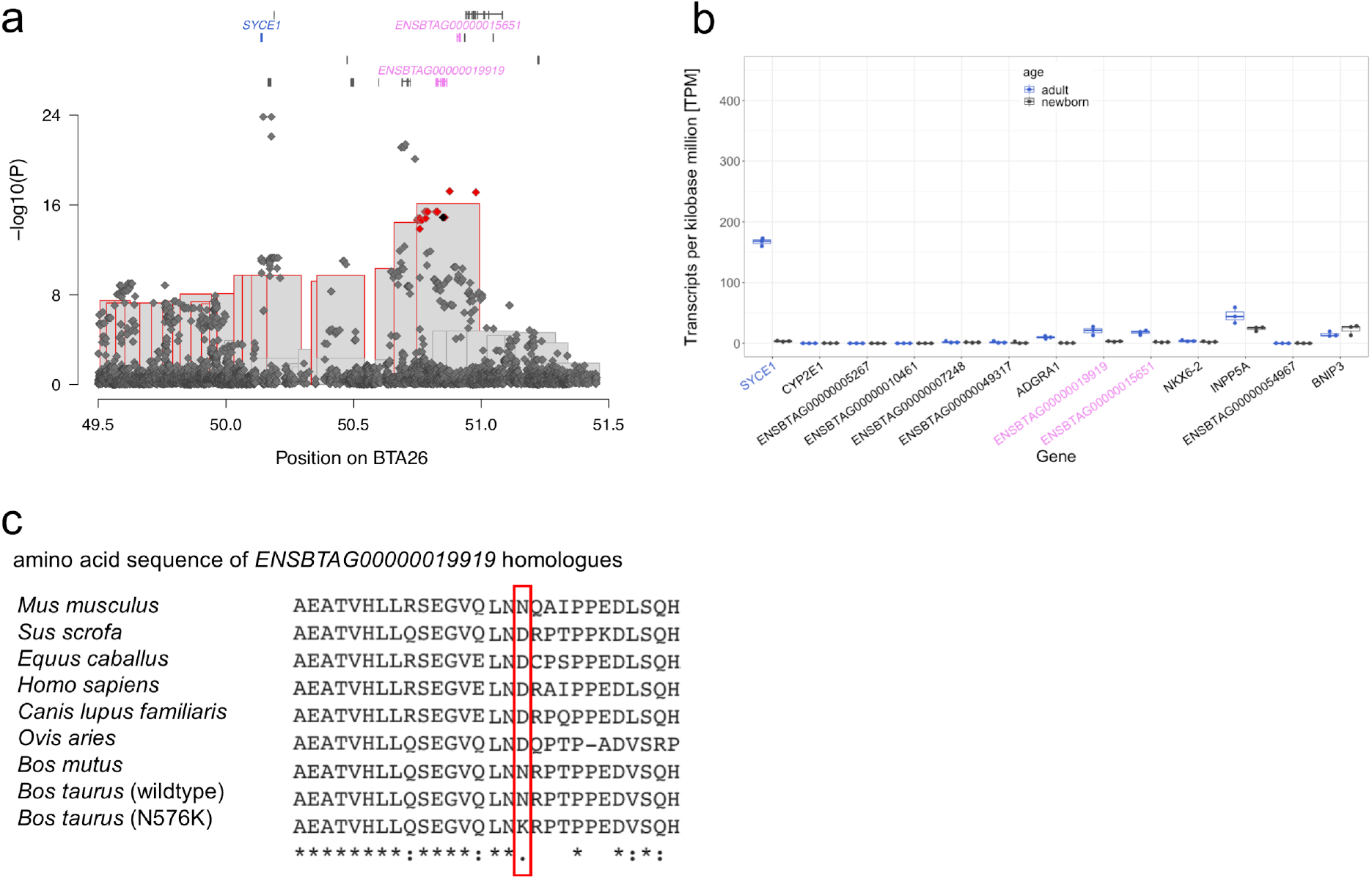
Fine mapping of variants associated with male fertility on BTA26. **a** Zoom on the Manhattan plot on chromosome 26, 49.5 to 51.5 Mb, of the whole-genome GWAS on bull fertility with recessive model. Bonferroni corrected significant variants are displayed in red. The *ENSBTAG00000019919* missense variant (BTA26:50,850,915C>G) is indicated with a black dot. Symbols over the Manhattan plot indicate genes and transcripts that are annotated at the QTL region. Blue and violet colour indicates genes and transcripts with testis-biased expression and male and female reproductive tract specific expression. **b** Expression of genes between 49.5 and 51.5 Mb in adult (blue) and newborn (black) bull testis in TPM. Blue and violet labels indicate genes with testis-biased expression and male and female reproductive tract specific expression. **c** Clustal Omega multi-species alignment of the protein produced from *ENSBTAG00000019919* and its homologous genes in *Mus musculus* (ENSMUST00000129990.8), *Sus scrofa* (ENSSSCT00000011782.4), *Equus caballus* (ENSECAT00000077103.1), *Homo sapiens* (ENST00000368586.10), *Canis lupus familiaris* (ENSCAFT00000017743.4), *Ovis aries* (ENSOART00000018738.1), *Bos mutus* (ENSBMUT00000020335.1), *Bos taurus* wildtype (ENSBTAT00000026536.5), and *Bos taurus* mutant (N576K).

The association study with imputed sequence variant genotypes revealed 18 variants that were stronger associated with bull fertility than the p.Asn576Lys-variant. The two variants with the lowest P value (P = 1.5 × 10^−24^) were BTA26:50145932C>T (rs720936782) and BTA26:50178052C>T (rs210673393), that are 4375 bp downstream of the stop codon of *SYCE1* encoding synaptonemal complex central element protein 1 and 2171 bp upstream of the start codon of *CYP2E1* encoding cytochrome P450, family 2, subfamily E, polypeptide (Figure 7). *SYCE1* is expressed at 167 TPM in adult bull testis, whereas *CYPE1* is not expressed (TPM < 1) (Figure 7b). However, both variants were not compatible with the inheritance of the top haplotype.

In order to disentangle the QTL, we performed haplotype and sequence-based association studies conditioned on either the top associated haplotype, the p.Asn576Lys-variant in *ENSBTAG00000019919* (BTA26:50,850,915C>G, rs378141069), or the BTA26:50145932C>T (rs720936782) variant downstream *SYCE1* (Supplementary Figure 5a,b,c). The conditional analyses indicated that the top haplotype and the p.Asn576Lys-variant capture the QTL variance only partially, because the rs720936782 variant was still associated with bull fertility (P = 1.1 × 10^−9^ and 8.5 × 10^−9^), albeit when conditioned on rs378141069 not at the stringent Bonferroni-corrected significance threshold (3.88 × 10^−9^). However, the QTL was absent when the analyses were conditioned on rs720936782 (Supplementary Figure 5c). In our set of the imputed sequence variant genotypes, rs378141069 and rs720936782 were in moderate linkage disequilibrium (r^2^ = 0.598).

### Variants compatible with the BTA18 QTL

The most significantly associated haplotype (P = 5.4 × 10^−9^) for bull fertility on BTA18 was between 36,256,440 and 36,555,965 bp. The haplotype was not associated with any semen quality or sperm morphology trait. The haplotype segregates in the BSW population at a frequency of 0.32. Of 3736 bulls from the mapping cohort, 434 were homozygous and 1690 heterozygous carriers of the top haplotype. Of 125 BSW bulls with whole-genome sequence variant genotypes, 14 were homozygous and 52 heterozygous haplotype carriers. Sequence variant genotyping revealed 30,018 polymorphic sites between 33,256,440 and 39,555,965 bp, of which 212 were compatible with the recessive inheritance of the top haplotype. Four compatible variants were located in protein-coding regions: a missense variant in *NFAT5* (BTA18:36,768,945, P = 1.5 × 10^−4^, SIFT score: 0.3 (tolerated)), and three synonymous variants in *ENSBTAG00000052086* (BTA18:36,734,722G>A, rs378665712, P = 1.5 x 10^−4^), *NQO1* (18:36789835C>T, rs110531779, P = 1.5 × 10^−4^) and *NOB1* (BTA18:36,816,759A>C, rs41874533, P = 1.5 × 10^−4^). *NFAT5*, *NQO1*, and *NOB1* are expressed at 22, 16, and 6 TPM in adult bull testes. Association testing between imputed sequence variant genotypes and bull fertility revealed four non-coding variants that exceeded the Bonferroni-corrected significance threshold (3.88 × 10^−9^, Figure 6a): a variant considered upstream *NIP7*, downstream *TMEMD6,* and intronic *COG8* (BTA18: 36,480,384T>G, rs380735496, P = 1.9 × 10^−9^), an intron variant in *TERF2* (BTA18:36,500,410A>C, rs378216493, P = 1.9 × 10^−9^), and two intronic variants in *CYB5B* (BTA18:36,561,644T>A and BTA18:36,565,464A>G, rs136784976 and rs379081158, both P = 3.3 × 10^−9^). These variants were as well compatible with the recessive inheritance of the top haplotype. *NIP7*, *TMED6*, *COG8*, *TERF2* and *CYB5B* are expressed at 38, 86, 35, 50, and 73 TPM in testis tissue of adult bulls (Figure 6b).

**Figure 6:**
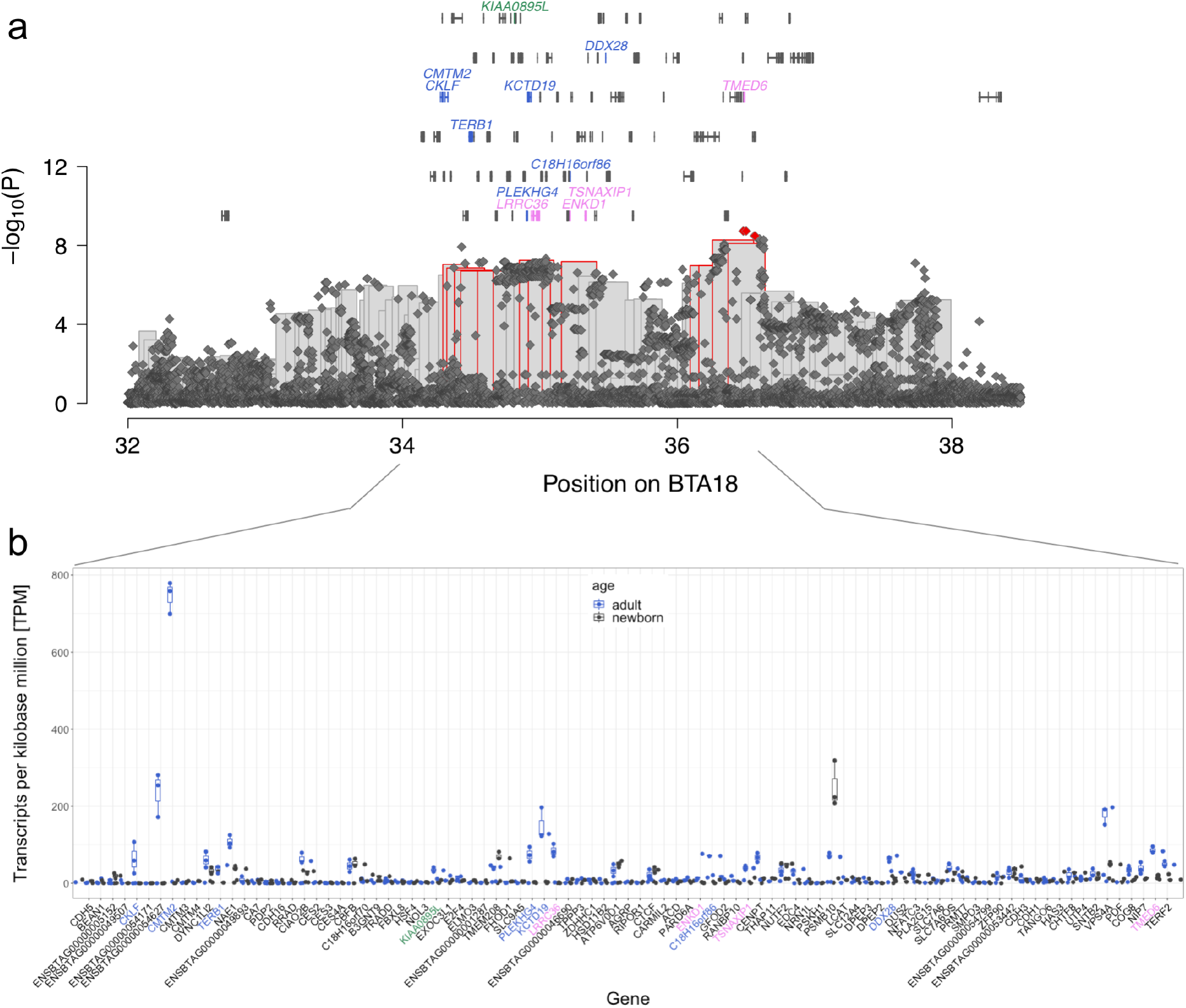
Detailed view of a QTL for bull fertility at BTA18. **a** Manhattan plot representing the association of imputed sequence variants (diamonds) and haplotypes (bars) located on chromosome 18 between 32 and 38.5 Mb. Red framed bars and diamonds represent haplotypes and variants that meet the Bonferroni-corrected significance threshold. Symbols over the Manhattan plot indicate genes and transcripts that are annotated at the QTL region. Blue, violet, and green colour indicates genes and transcripts with testis-biased expression, male and female reproductive tract specific expression, and testis and brain specific expression. **b** Expression of genes within the segment 34 to 36.5 Mb in adult (blue) and newborn (black) bull testis in TPM. Blue, violet, and green labels indicate genes with testis-biased expression, male and female reproductive tract specific expression, and testis and brain specific expression.

## Discussion

In a mapping cohort of 3736 BSW bulls from which semen was collected at semen collection centers under standardized conditions, only 10 % of the phenotypic variation of bull fertility was explained by autosomal variants. This estimate suggests that the additive genetic heritability of bull fertility is low in BSW cattle, which agrees with findings in other cattle populations ^19–23^. It is possible that we underestimated the genomic heritability of bull fertility, because the genomic relationship matrix was constructed from autosomal SNPs only. Variants on the sex chromosomes may also contribute to inherited variation in bull fertility ^24,25^. Our mapping cohort provided sufficient statistical power to detect large-effect QTL for a complex trait with an estimated narrow-sense heritability of 10 % ^26^. However, we did not detect additive QTL. Therefore, we conclude that the additive genetic architecture of bull fertility is polygenic and determined by QTL with small to moderate effects.

Our haplotype-based association analysis identified five recessive QTL with moderate frequencies that, taken together, had a substantial effect on bull fertility in the BSW cattle population. Homozygous haplotype carriers are fertile, but their fertilization success is reduced in artificial inseminations. Our data did not allow investigating bull fertility in natural matings. Phenotypic manifestations of the five recessive QTL detected in our study are less detrimental than effects arising from pathogenic alleles in *CCDC189*, *ARMC3* and *TMEM95* that lead to male sterility in cattle ^27–29^. Yet, the hypothetical exclusion of bulls carrying at least one of the five detected top haplotypes in the homozygous state from breeding would significantly increase the 56-day non-return rate in cows and heifers. Thus, the recessive QTL detected in our study contribute to quantitative variation in bull fertility. Similar findings in Fleckvieh, Jersey, and Holstein bulls suggest that the genetic architecture of male reproductive traits is shaped by non-additive effects ^8,11,29^. Repeatability estimates for traits describing male reproductive performance in cattle are substantially higher than narrow-sense heritability estimates in different cattle breeds including Brown Swiss ^13,30,31^. In addition to additive effects, the repeatability also includes permanent environmental and non-additive sources of variation. Our study shows that non-additive QTL for bull fertility can be detected in large mapping cohorts using genome-wide association testing. The fact that non-additive effects contribute substantially to the genetic variation of male reproductive traits has implications in efforts to predict bull fertility from dense molecular markers. While improving the fertility of bulls is not a breeding objective in most cattle improvement programs, the genome-based prediction of bull fertility might facilitate implementing compensation strategies for bulls for which the anticipated insemination success is low. Statistical models that consider only additive effects have low predictive power for bull fertility ^8,32^. Our findings suggest that non-additive effects should be considered in order to improve the accuracy of genomic prediction for male reproductive traits.

The sequence-based fine-mapping of the QTL revealed candidate causal variants in coding sequences of genes that are either testis-specific expressed in cattle or contribute to variation of male fertility in species other than cattle. The top haplotype on BTA6 is in strong linkage disequilibrium with a likely causal variant that activates cryptic splicing in *WDR19* ^13^. A variant in human *WDR19* is associated with morphological abnormalities of sperm and impaired male fertility ^33^. The top haplotypes at the BTA1, BTA25 and BTA26 QTL encompass putatively deleterious variants in genes (*SPATA16*, *VWA3A, ENSBTAG00000006717* and *ENSBTAG00000019919*) that are expressed in the bovine male reproductive tract. Furthermore, *SPATA16* and *ENSBTAG00000006717* are specifically expressed in the male reproductive tract (http://cattlegeneatlas.roslin.ed.ac.uk) and the produced proteins SPATA16 and ABCA14-like are prevalent in spermatozoa ^34,35^. Pathogenic variants in *SPATA16* and *ENSBTAG00000019919* orthologs have been associated with either impaired male fertility in humans or impaired movement of flagella in a flagellated green alga ^18,36^. VWA3A is associated with motile cilia in humans ^17^. Candidate causal variants for recessive traits are frequently prioritized if they are compatible with the segregation of the top haplotype ^13,37–39^. Strictly applying this criterion, the missense and nonsense variants in the coding sequences of *SPATA16*, *VW3A3, ENSBTAG00000006717* and *ENSBTAG00000019919* qualify as compelling candidate causal variants for impaired bull fertility; these variants are in linkage disequilibrium with the respective top haplotype and predicted to be deleterious to protein function. Our screen for variants that are compatible with the inheritance of the top haplotype takes genotyping errors into account. However, due to the low number of sequenced bulls that were homozygous haplotype carriers, it is possible that some variants were misclassified. Thus, we performed a sequence based association study as a complementary fine-mapping approach. An association study with imputed sequence variant genotypes showed that the candidate causal variants are also associated with bull fertility to a certain degree. Yet, non-coding variants were stronger associated with bull fertility than the putatively functional coding variants at all QTL. It is possible that the true causal variants reside in regulatory regions, but are in linkage disequilibrium with the coding variants prioritized in our study. Such a pattern could also indicate multiple trait-associated alleles within a QTL ^40^.

The p.Asn576Lys variant in *ENSBTAG00000019919* is a particularly compelling candidate causal variant for the BTA26 QTL because it was (i) in linkage disequilibrium with the top haplotype, (ii) highly significantly associated (P=1.3 × 10^−15^) in the sequence based association study, (iii) a lysine at position 576 is predicted to be deleterious to protein function, and (iv) the expression of *ENSBTAG00000019919* is restricted to the male and female reproductive tract. However, the sequence-based association study also revealed highly significantly associated variants nearby *SYCE1* that were not in complete linkage disequilibrium with the top haplotype. In fact, an association analysis conditional on the p.Asn576Lys variant did not fully account for the BTA26 QTL variation. The non-coding variant downstream *SYCE1* captured the QTL variance better. *SYCE1* is also a candidate gene for imapired male fertility, because it is testis-specific expressed at very high levels (TPM > 150). Loss of function of SYCE1 leads to spermatogenic arrest in human and mice ^41–43^. The Synaptonemal Complex Central Element Protein 1 is part of the central element of the synaptonemal complex that is required for chromosome synapsis in prophase I of meiosis ^44^. Errors in the meiotic recombination can lead to sperm aneuploidy ^45^ and thus to embryonic loss ^46,47^. It seems plausible that sequence variation affecting *SYCE1* and the p.Asn576Lys variant in *ENSBTAG00000019919* contribute to the BTA26 QTL variation.

It is worth mentioning, that the true causal variants may not necessarily be the most significantly associated variants in association studies with imputed sequence variant genotypes due to imputation errors or sampling effects ^48–50^. Interestingly, also the BTA6:58373887C>T variant (rs474302732) that had been reported as a putatively causal variant for the BTA6 QTL ^13^, was not the top variant in our association study between imputed sequence variant genotypes and bull fertility. Thus, we consider the coding variants in *SPATA16*, *VWA3A, ENSBTAG00000006717* and *ENSBTAG00000019919* as candidate causal variants for impaired male reproductive performance, but further functional investigations and replication studies in independent populations are required to corroborate their association with bull fertility.

Because our association studies considered both bull fertility and semen quality, we were able to reveal the mechanisms through which the BTA1 QTL contributes to variation in fertility. Bulls that are homozygous for the top haplotype encompassing *SPATA16* are less fertile likely because their ejaculates contain an increased number of sperm with morphological abnormalities of the sperm head. Abnormal head morphology is negatively correlated with NRR56 ^51,52^. However, semen quality of homozygous bulls is not as severely compromised as, e.g., in males with globozoospermia ^36,53^ and spermatogenic arrest ^54^, respectively, due to loss-of-function alleles in human and murine *SPATA16*. It is plausible that the SPATA16:p.Ile193Met variant compromises semen quality in the homozygous state, as the methionine at position 193 is predicted to be deleterious to SPATA16 function. However, we can not preclude that non-coding variants in linkage disequilibrium with p.Ile193Met contribute to the BTA1 QTL variation. In any case, the isoleucine residue at position 193 of SPATA16 is not essential for fertilization, because homozygous bulls are fertile. Sperm head shape abnormalities were only detected in fixed and therefore immotile spermatozoa of homozygous haplotype carriers. Because the morphological abnormalities of the sperm heads are subtle, they remain unnoticed in fresh and motile spermatozoa. In fact, the majority of sperm produced from homozygous haplotype carriers is normal, but the overall proportion of normal sperm is reduced. Many bulls homozygous for the BTA1 QTL are judged to be not suitable for breeding due to insufficient semen quality before their semen is used for artificial inseminations. Thus, the actual effect of the BTA1 QTL on bull fertility is likely more pronounced than observed in our study that was based on insemination success from ejaculates meeting minimum quality requirements.

We found no clues from the analysis of semen quality data why fertility is reduced in bulls that are homozygous carriers of the top haplotypes at the BTA18, BTA25, and BTA26 QTL. Thus, our study confirms indirectly that routine semen quality parameters, once bulls producing ejaculates with clearly insufficient quality are removed, are not sufficient to anticipate bull fertility ^55^. Computer-assisted and flow-cytometric sperm analyses may provide further insights into mechanisms that underpin phenotypic variation in bull fertility ^56^. Moreover, the phenotype bull fertility does not enable differentiating between fertilization failure and early embryonic losses, because it is based on the 56-day non-return rate. It is thus possible that fertilization is normal, but the apparently reduced bull fertility is primarily a manifestation of early embryonic losses ^57,58^.

## Methods

### Genotypes of BSW bulls

Microarray-derived genotypes of 33,045 BSW cattle were provided by breeding associations. The genotypes were determined using various versions of Illumina BovineSNP Bead chips comprising between 20K and 777K SNP. Quality control including the removal of variants and samples with more than 10 % missing genotypes and the exclusion of variants deviating significantly (P < 10^−8^) from Hardy-Weinberg proportions was carried out using *plink* (v1.9) ^59^.

Using a reference panel of 1166 cattle that had Illumina BovineHD-derived genotypes for 777,962 SNPs, the genotypes for animals typed at lower density were imputed to higher density using *Beagle 5.1* ^60^ with default settings. The final dataset comprised 33,045 animals and 683,609 SNPs. For association testing, we considered only 3881 bulls for which either bull fertility or semen quality data were available.

### Bull fertility in Swiss BSW sires

Bull fertility (as at February 2020) was provided by Swissgenetics for 1382 BSW bulls that were used for between 232 and 15,690 first inseminations with conventional (i.e., semen was not sorted for sex) frozen-thawed semen. Bull fertility was estimated based on the 56-day non-return rate after the first insemination in cows and heifers using a linear mixed model similar to the one proposed by Schaeffer et al. ^2^: *y*_*ijklmnopq*_ = μ + *MO*_*i*_ + *PA*_*j*_ + *PR*_*k*_ + (*BS* × *BK*)_*lm*_ + *TE*_*n*_ + *h*_*o*_ + *s*_*lp*_ + *e*_*ijklmnopq*_, where **y**_ijklmnopq_ is either 0 or 1, indicating whether or not a subsequent insemination was recorded within a 56-day interval after the first insemination, μ is the intercept, **MO**_i_ is the insemination month, **PA**_j_ is the parity (heifer or cow), **PR**_k_ is the cost of the semen dose, (**BS** × **BK**)_lm_ is a combination of bull’s breed x cow’s breed, **TE**_n_ is the insemination technician, ho is the random effect of the herd, **s**_lp_ is the random effect of the bull expressed as deviation from the average non-return rate, and **e**_ijklmnopq_ is a random residual term. Bull fertility was subsequently standardized to a mean of 100 ± 12. The resulting value is an objective measurement of bull fertility. Three phenotypic outliers (i.e., bulls for which the fertility was more than 5 standard deviations below average) were not considered for subsequent analyses because such bulls might carry rare genetic conditions ^29^ that can lead to spurious associations ^61^. For the GWAS, we considered 1130 Swiss BSW bulls for which we also had partially imputed genotypes. Their fertility was quantified based on 850,708 first artificial inseminations.

### Bull fertility in German and Austrian BSW sires

Phenotypes for bull fertility were provided by ZuchtData EDV-Dienstleistungen GmbH, Austria, for 4617 BSW bulls from Germany and Austria (as at December 2017) that were used for 4,267,990 and 1,646,254 inseminations in cows and heifers, respectively. Bull fertility was obtained from routine breeding value estimation for reproductive traits, which are jointly estimated for males and females ^62^. The statistical model used to estimate breeding values for female reproductive traits includes a fixed effect for the service sire that represents bull fertility as deviation from the population mean. Bull fertility was subsequently standardized to a mean of 100 ± 12. We excluded four bulls with exceptionally poor fertility (see above). For association testing, we considered 2606 bulls from the German and Austrian BSW populations for which we also had partially imputed genotypes.

### Compilation of a mapping cohort for association testing of bull fertility

To estimate the genetic correlation of bull fertility between the Swiss and German / Austrian bulls, a bivariate restricted maximum likelihood (REML) analysis was performed using the *GCTA* software (version 1.93.2) ^63^. We fitted a genomic relationship matrix that was built using 589,791 autosomal SNPs with minor allele frequency greater than 0.5 %. To take population stratification into account, we fitted the top 10 principal components of the genomic relationship matrix. The REML estimation revealed that the same genetic determinants control the trait in both mapping cohorts (rg=1 ± 0.28). We standardized the phenotypes within the Swiss and German / Austrian cohorts to compile a joint mapping population of 3736 bulls. The proportion of phenotypic variation explained by 589,791 autosomal SNPs with minor allele frequency greater than 0.5 % was estimated using *GCTA*.

### Semen quality of Swiss BSW bulls from routine fresh semen assessment

Semen quality data were provided by Swissgenetics for 74,945 ejaculates that were collected from 1432 BSW bulls between January 2000 and November 2019. Parameters assessed for fresh ejaculates were semen volume (in ml), sperm concentration (million sperm per ml) quantified using photometric analysis, and sperm motility (percentage of sperm with forward motility) assessed visually using a heated-stage microscope at 200-fold magnification. Moreover, the amount of sperm with head and tail anomalies was classified for each ejaculate with scores ranging from 0 to 3 (0: no or very few anomalies, 1: less than 10 % sperm with anomalies, 2: between 10 and 30 % sperm with anomalies, 3: more than 30 % sperm with anomalies). Ejaculates that fulfilled the requirements for artificial insemination (semen volume above 1 ml, more than 300 million sperm per ml, at least 70 % motile sperm, no apparent impurities and no excessive amount of sperm with head and tail abnormalities) were diluted using a Tris-egg yolk based extender, filled in straws containing between 15 and 25 million sperm, and cryoconserved in liquid nitrogen. The raw ejaculate data were filtered according to the parameters listed in Table 2. Response variables for the association study were the mean semen quality values of 902 bulls assessed in at least eight ejaculates per bull.

**Table 2:** Parameters and values applied to filter the semen quality data of Swiss BSW bulls.

| Filtering parameter | Number of bulls | Number of ejaculates |
| --- | --- | --- |
| Raw data | 1432 | 74,945 |
| Age at ejaculate collection > 400 and < 1000 days | 1380 | 60,246 |
| Interval to preceding ejaculate known (without first ejaculate) | 1379 | 59,353 |
| Ejaculate volume recorded | 1366 | 58,846 |
| No mixed ejaculates | 1358 | 42,870 |
| Only processed ejaculates | 1311 | 38,370 |
| Interval between ejaculates <= 8 days | 1304 | 38,023 |
| Only first ejaculate per day | 1301 | 34,665 |
| Sperm motility >= 70 % | 1285 | 34,510 |
| Anomaly score plausible | 1285 | 34,505 |
| Sperm per straw >= 15 million (not sorted for sex) | 1285 | 34,278 |
| Semen collector and bull handler known | 1253 | 32,023 |
| (imputed) HD genotype available | 1119 | 29,814 |
| >= 8 records per bull | 902 | 28,856 |

### Sperm morphology data of Swiss BSW bulls from breeding soundness examinations

Sperm morphology was provided by Swissgenetics for 1766 ejaculates of 669 BSW bulls that entered the semen collection center between August 2009 and February 2020. Technicians systematically evaluate sperm morphology of prospective breeding bulls as part of the andrological examination in formalin-fixed fresh semen samples under oil immersion light microscope. Between 150 and 200 sperm per ejaculate are classified according to three non-compensatory and 13 compensatory sperm morphology categories that are considered as either minor or major defects (Table 3). For ejaculates that evidently fail the quality assessment, only 50 spermatozoa are classified. For sperm with multiple morphological abnormalities, only the most severe defect is recorded. Bulls producing ejaculates that contain at least 75 % normal sperm and not more than 20 % sperm with non-compensatory defects are considered suitable for breeding. If these requirements are not fulfilled, the sperm morphology examination is repeated approximately four weeks later. If a bull produces ejaculates that contain between 65 and 74 % normal sperm and not more than 20 % sperm with non-compensatory defects, its ejaculates are processed but the reduced semen quality is compensated with more sperm per insemination dose. Bulls that repeatedly produce ejaculates with less than 65 % normal sperm and more than 20 % sperm with non-compensatory defects are not used for breeding.

**Table 3:** Sperm morphology traits examined in Swiss BSW bulls. Description of sperm morphology traits and their classification into compensatory or non-compensatory and major or minor defects as used in the breeding soundness examination of bulls at the Swiss artificial insemination centre Swissgenetics.

| Defect number | Description | Compensatory or non-compensatory | Major or minor |
| --- | --- | --- | --- |
| 0 | Abnormal head shape (pyriform / pear shape, tapered / narrow head base, others) | Non-compensatory | Major |
| 1 | Vacuoles | Non-compensatory | Major |
| 2 | Condensed DNA | Non-compensatory | Major |
| 3 | Micro- or macroencephaly | Compensatory | Minor |
| 4 | Loose heads | Compensatory | Minor |
| 5 | Knobbed acrosome | Compensatory | Major |
| 6 | Detached acrosome | Compensatory | Minor |
| 7 | Dag defect | Compensatory | Major |
| 8 | Curled tail (clef-like) | Compensatory | Major |
| 9 | Middle-piece knobbed | Compensatory | Major |
| 10 | Tail abaxially attached to head | Compensatory | Minor |
| 11 | Tail end loop | Compensatory | Minor |
| 12 | Cytoplasmic droplet proximal (middle-piece near head) | Compensatory | Major |
| 13 | Cytoplasmic droplet distal (middle-piece near tail) | Compensatory | Minor |
| 14 | Underdeveloped | Compensatory | Major |
| 15 | Double form | Compensatory | Minor |

For the association study, we considered three trait categories of the sperm morphology examination:

- The total number of examinations per bull (n = 595 bulls with 1577 ejaculates). The number of examined ejaculates ranged from 1 to 26 (median 2).
- The average proportion of sperm for 13 defect categories (Table 3) was quantified for 575 bulls based on 1165 examined ejaculates for which more than 150 spermatozoa were assessed. Between 1 and 13 (median=1) ejaculates per bull were examined.
- Aggregate phenotypes were the average proportion of sperm with normal morphology (assessed sperm - sum of all morphological defects), major defects, minor defects, compensatory defects and non-compensatory defects (Table 3).

### Haplotype-based association testing

Haplotype-based association testing was implemented in R using a sliding window-based approach that was previously applied to map complex and binary traits ^13,64^. We shifted a window of 50 contiguous SNPs in steps of 15 SNPs along the autosomal haplotypes. Within each sliding window, all haplotypes with a frequency above 1 % were tested for association with bull fertility or semen quality using the linear model 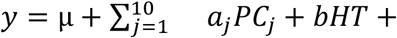 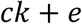, where y is a vector of phenotypes (see above), μ is the intercept, **PC**_j_ are the top 10 principal components of the genomic relationship matrix (see above), **a**, **b** and **c** are effects of the principal components, the haplotype (**HT**) tested and a covariable **k** (e.g., the recessive BTA6 haplotype associated with male fertility), respectively, and **e** is a vector of residuals that are assumed to be normally distributed. Haplotypes were tested for association assuming either additive or recessive mode of inheritance. Recessive tests were carried out for haplotypes that were observed in the homozygous state in at least 0.5 % of individuals. The genomic inflation factor lambda was calculated as: lambda = median(qchisq(1-p, 1)) / qchisq(0.5, 1) where p is a vector of P values.

### Redefining a previously detected fertility-associated haplotype on BTA6

We previously detected a recessive QTL for semen quality and bull fertility on BTA6 ^13^ that segregates at high frequency in BSW cattle. In order to localize the QTL in our mapping cohort, we performed an association study between bull fertility and 6962 haplotypes (31,362 partially imputed SNPs) of chromosome 6 in 1130 Swiss BSW bulls using a haplotype-based association mapping approach (see above) assuming recessive inheritance. The association study revealed the strongest association (P = 2.4 × 10^−20^) with bull fertility for a haplotype located between 57,628,627 and 57,919,676 bp. In association analyses that were conditioned on the BTA6 QTL, we fixed the bulls’ statuses (recessively coded: non-carrier and heterozygous vs. homozygous) for this haplotype as a covariate.

### Whole-genome sequencing, sequence variant genotyping and imputation

We considered 368 BSW and Original Braunvieh (OBV) cattle that were sequenced at between 3 and 60-fold (median of 11.6) genome coverage using either Illumina HiSeq 2500, Illumina HiSeq 4000, or Illumina NovaSeq6000 instruments. The raw sequencing reads were filtered using the *fastp* software ^65^ with default parameter settings and subsequently aligned to the ARS-UCD1.2 assembly of the bovine genome ^66^ using the mem-algorithm of the *Burrows Wheeler Aligner (BWA)* software package ^67^. Single nucleotide and short insertion and deletion polymorphisms were detected and genotyped using the *HaplotypeCaller*, *GenomicsDBImport* and *GenotypeGVCF* modules of *GATK* ^68^. Low quality variants were filtered out using *GATK*’s best practice recommendations for hard filtering. Following filtration, 24,614,962 SNPs and 3,981,732 indels were retained for subsequent analyses. *Beagle 4.1* ^69^ phasing and imputation was applied to improve the primary genotype calls from *GATK* and infer missing genotypes. We used the *Beagle 5.1* ^60^ software with default settings to impute the genotypes for 28,596,694 sequence variants for 3736 bulls from the mapping cohort that had partially imputed genotypes at 683,609 SNPs from a reference panel of 368 sequenced BSW and OBV animals.

### Sequence-based association study

Imputed sequence variant genotypes that had allele frequency greater than 0.05 and imputation accuracy (*Beagle* r^2^) greater than 0.4 were considered for association testing. The statistical model applied to test for an association between the imputed sequence variants and bull fertility was analogous to the haplotype-based association study. We used the *plink* (version 1.9) ^59^ software to fit a linear regression model assuming a recessive mode of inheritance conditional on the top ten principal components of a genomic relationship matrix (see above) and the fertility-associated haplotype on BTA6 (see above).

### Prioritization of candidate causal variants

Variants that exceeded the Bonferroni-corrected significance threshold of 3.88 × 10^−9^> (defined based on 12,861,528 imputed sequence variants with MAF > 0.05 and *Beagle* r^2^ > 0.40) and were compatible with the inheritance of the top haplotype were considered as candidate causal variants. To identify variants compatible with the inheritance of the top haplotype, we considered sequence variant genotypes from 125 BSW cattle with bull fertility phenotypes that had both array-derived genotypes and sequence variant genotypes. Their carrier status for the top haplotype at the QTL was inferred from the array-derived genotypes. We considered sequence variants within ± 3 Mb of the top haplotype as positional candidate causal variants. We filtered for variants that had allele frequency

- greater than 0.8 in animals that carried the top haplotype in the homozygous state,
- between 0.4 and 0.6 in heterozygous haplotype carriers,
- and less than 0.05 in non-carrier animals.

Functional consequences of candidate causal variants were predicted based on the Ensembl (release 101, August 2020) annotation of the ARS-UCD1.2 *Bos taurus* assembly using the *Variant Effect Predictor* (VEP) software tool ^70^.

### Bioinformatic analyses

The *Clustal Omega* ^71^ software was used for multi-species alignment of protein sequences and *Missense3D* ^72^ to predict putative effects of missense variants on protein tertiary structure. The three dimensional protein structure was modelled using the *SWISS-MODEL* platform ^73^.

### Transcriptome analysis

To assess the abundance of transcripts in bovine testis, we downloaded between 47 and 58 million 2 x 150 bp paired-end sequencing reads from the European Nucleotide Archive (ENA, https://www.ebi.ac.uk/ena) that were generated using RNA extracted from testis tissue samples of three mature bulls and three newborn male calves of the Angus beef cattle breed (ENA accession numbers: SAMN09205186-SAMN09205191; ^74^). The RNA sequencing reads were pseudo-aligned to an index of the bovine transcriptome (ftp://ftp.ensembl.org/pub/release-98/fasta/bos_taurus/cdna/Bos_taurus.ARS-UCD1.2.cdna.all.fa.gz) and transcript abundance was quantified using the *kallisto* software ^75^. We used the R package *tximport* ^76^ to aggregate transcript abundances to the gene level.

### Imaging of spermatozoa from bulls homozygous for the haplotype on BTA1

Fresh semen samples were provided by Swissgenetics for two bulls that were homozygous carriers of the fertility-associated haplotype on BTA1. The haplotype status of the bulls was derived using microarray-derived genotypes. Sperm morphology was assessed in semen samples fixed in buffered formaldehyde saline solution using oil-immersion phase contrast microscopy (Olympus BX50, UplanF1 100×/1.30, Olympus, Wallisellen, Switzerland).

## Declarations

### Ethics declarations

#### Ethics approval

No ethics approval was required for the study. Semen samples of bulls were provided from an artificial insemination center after regular collection as part of their insemination straw production. Semen quality, sperm morphology, and insemination success were recorded at the Swiss artificial insemination center Swissgenetics to monitor bull fertility. Fertility records and genotypes were provided by breeding organizations and had been collected as part of normal breeding procedures. No samples were collected specifically for the present study.

## Competing interests

UW and FSH are employees of Swissgenetics. All other authors declare no competing interests.

## Acknowledgements

We highly appreciate the detailed explanations of sperm morphological examination at Swissgenetics by Susanne Meese and Sarah Wyck. We thank Sarah Wyck and Tjasa Kampara for providing semen for morphological analysis. We acknowledge Swissgenetics for providing phenotype data for the Swiss BSW bulls. We thank Franz Seefried for the identification of homozygous haplotype carriers in the Swiss BSW population. We thank Braunvieh Schweiz for providing genotype data of Swiss BSW bulls. We acknowledge the Arbeitsgemeinschaft Deutsches Braunvieh, Braunvieh Austria, Tierzuchtforschung Grub, the Chair of Animal Breeding of TU München, the Institute of Animal Breeding from Bayerische Landesanstalt für Landwirtschaft and ZuchtData EDV Dienstleistungen GmbH for providing genotype and phenotype data for the Austrian and German BSW bulls. We thank the Functional Genomics Center Zurich for generating DNA sequencing data. This study was financially supported from Swissgenetics, Zollikofen, Switzerland (https://swissgenetics.ch/) Grant-ID: 13453, the Förderverein Biotechnologieforschung e.V. (FBF), Bonn, Germany (https://www.fbf-forschung.de/) Grant-ID: 2-70024-18, and the Swiss National Science Foundation (310030_185229).

## Data availability

Whole-genome sequence data from 125 BSW cattle are available at the sequence read archive of the European Nucleotide Archive (https://www.ebi.ac.uk/ena), identifiers are given in Supplementary File 1. The results from 29 haplotype-based association studies as well as results from association studies between imputed sequence variants and bull fertility on Chromosomes 1, 18, 25, and 26 are available as Supplementary Data.

## Author information

### Affiliations

Animal Genomics, Institute of Agricultural Sciences, ETH Zürich, Eschikon 27, 8315 Lindau, Switzerland

Maya Hiltpold, Naveen Kumar Kadri, Hubert Pausch

Clinic of Reproductive Medicine, Vetsuisse Faculty, University of Zurich, 8057 Zurich, Switzerland Fredi Janett

Swissgenetics, 3052 Zollikofen, Switzerland Ulrich Witschi, Fritz Schmitz-Hsu

### Contributions

MH, NKK and HP analysed and interpreted the data. UW coordinated the collection of semen quality and sperm morphology data. FSH estimated bull fertility for the Swiss BSW bulls. FJ examined and imaged spermatozoa in phase-contrast light microscopy. MH and HP wrote the paper. All authors reviewed and approved the final version of the manuscript.

