## Supplementary Information for "Autosomal recessive loci contribute significantly to quantitative variation of male fertility in a dairy cattle population"

### Supplementary Figures

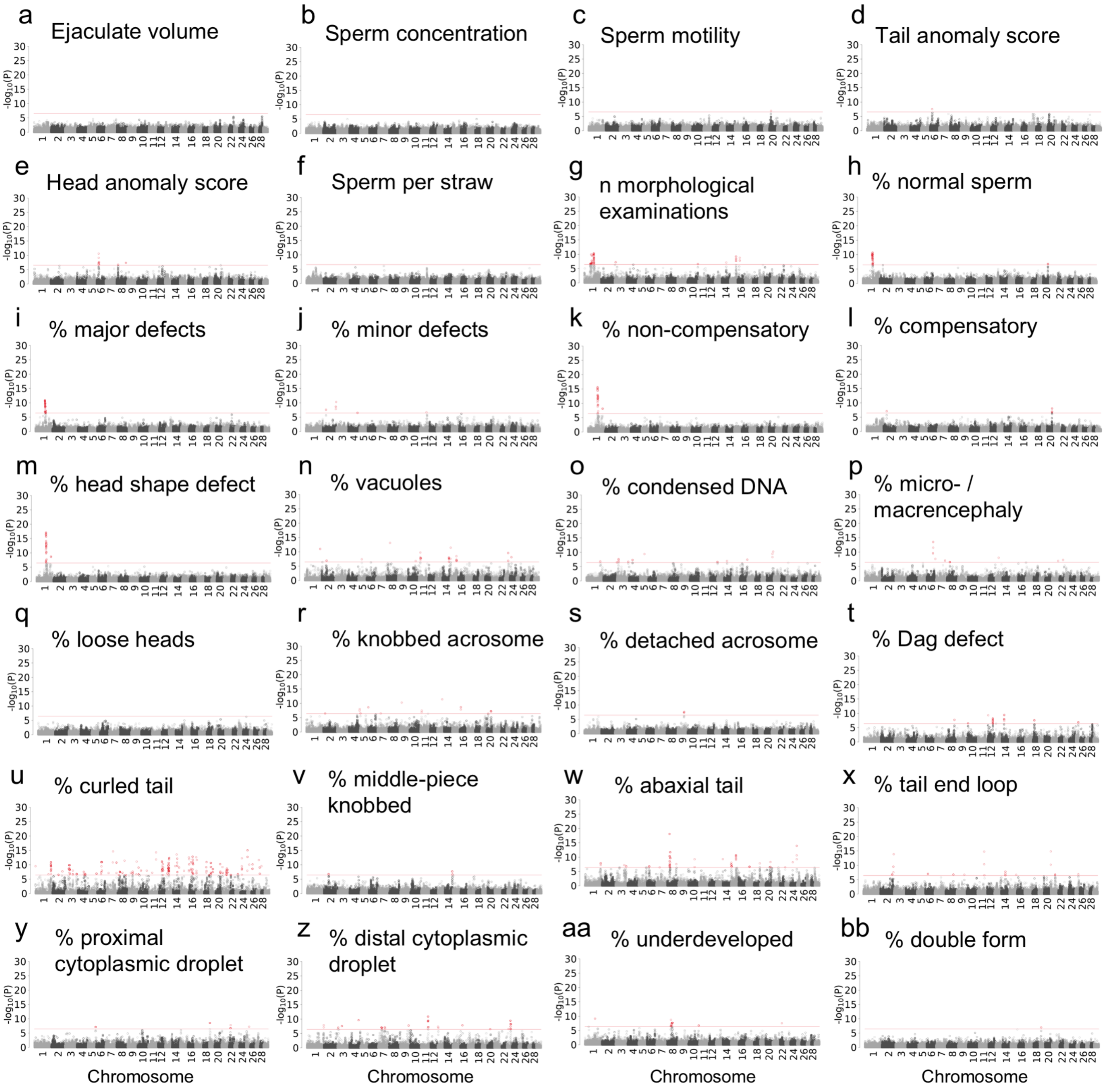


**Supplementary Figure 1: Haplotype-based genome wide association studies on bull fertility related traits.** Manhattan plots representing the association ($-\log_{10}(P)$) of haplotypes with 28 semen quality traits. The association tests were based on a recessive model and conditional on the top haplotype of the BTA6 QTL. Variants that exceed the Bonferroni-corrected significance threshold are displayed in red.

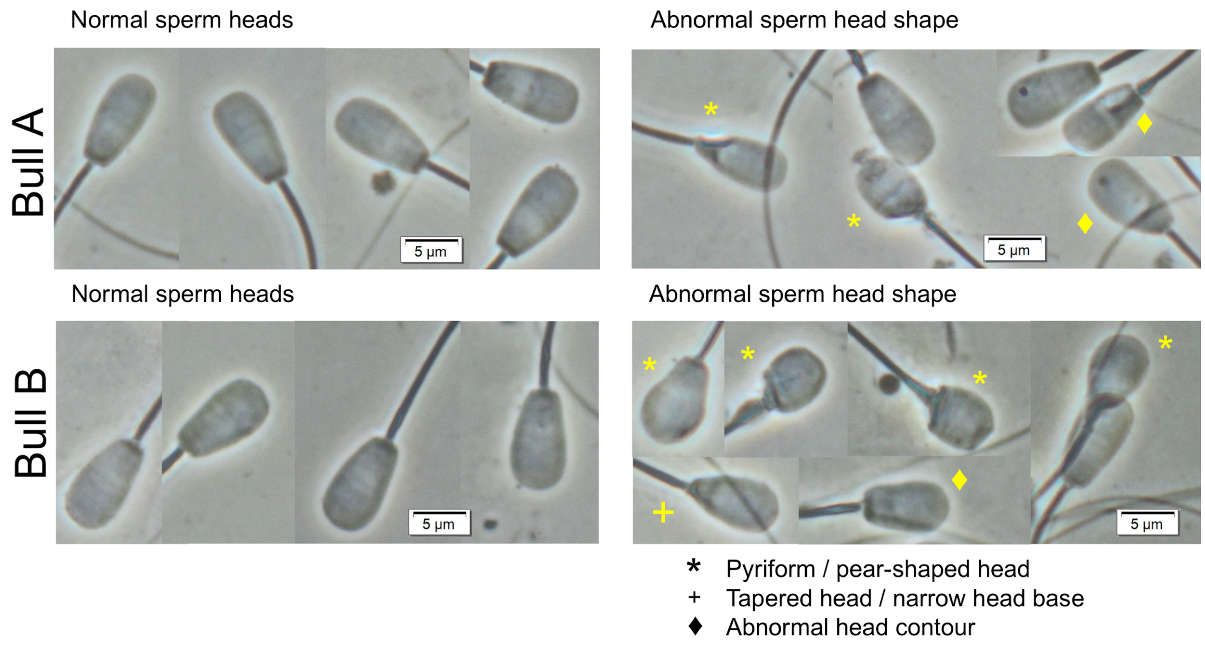


**Supplementary Figure 2: Bulls homozygous for the BTA1 haplotype produce sperm with head shape anomalies.** Detail view on phase-contrast microscopy images of spermatozoa two bulls homozygous for the BTA1 haplotype reveal both spermatozoa with normal morphology and spermatozoa with multiple morphological abnormalities of the head. The abnormal shapes consist of heads with rounder frontal part (pyriform / pear shape), narrower head base (tapered heads), and uneven shaped heads (abnormal head contour).

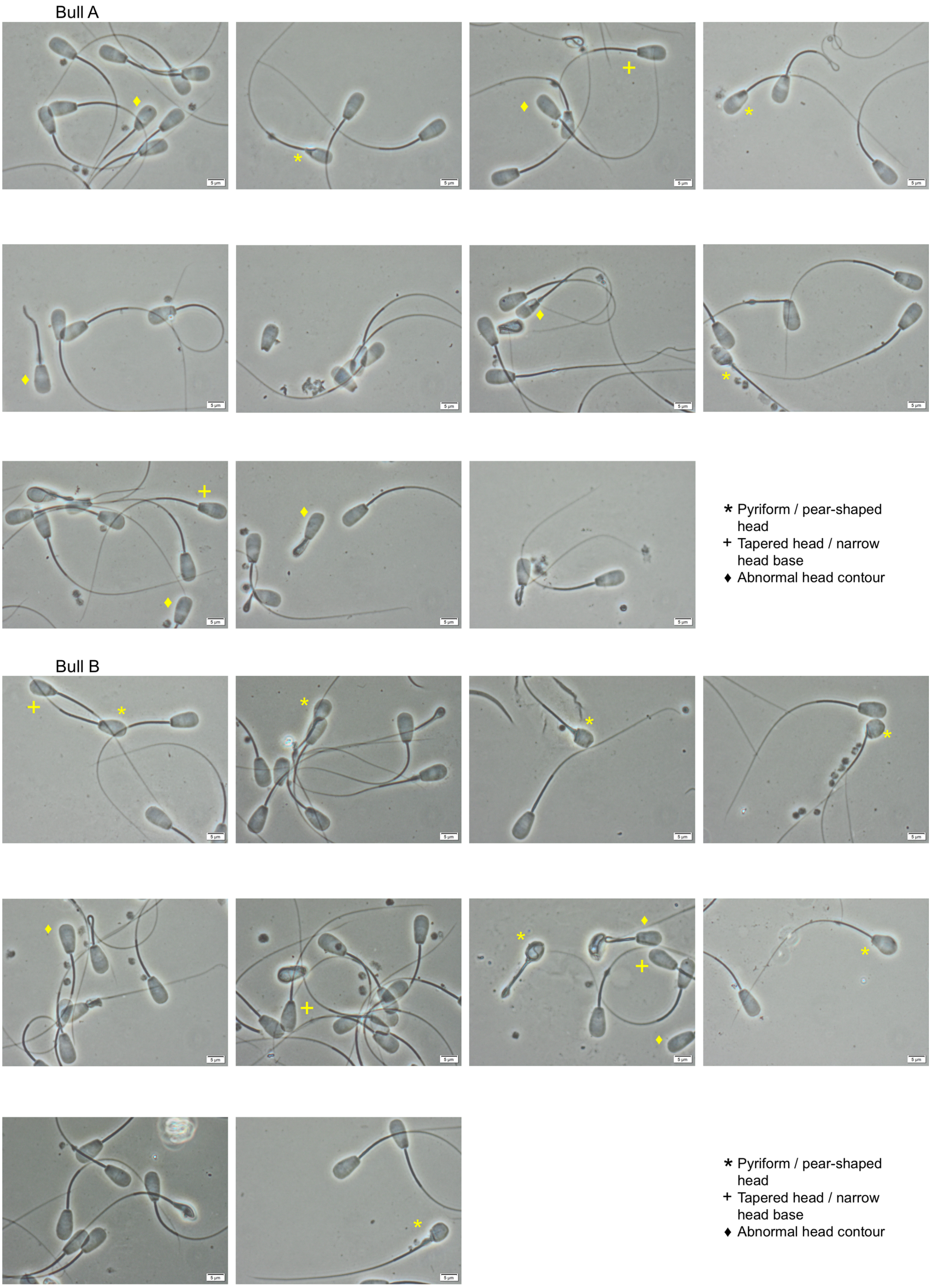


**Supplementary Figure 3: Sperm microscopy images of two bulls homozygous for the QTL on BTA1.** Complete phase-contrast microscopy images of detail view shown in Supplementary Figure 2.

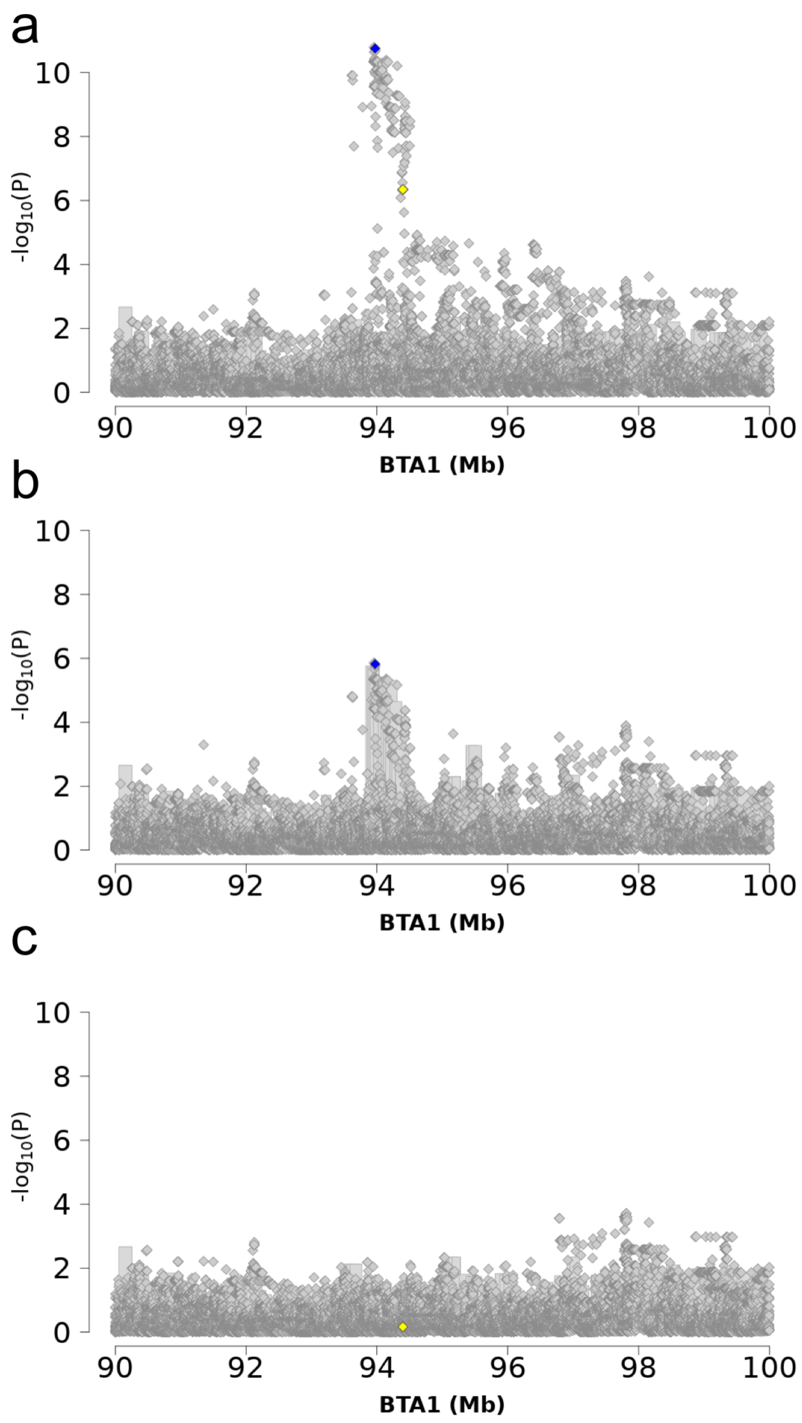


**Supplementary Figure 4: Conditional association analyses on BTA1 90-100 Mb. a** The imputed sequence variants (diamonds) and haplotypes (bars) association analyses were conditioned on the top associated haplotype on BTA1, **b** on the SPATA16:p.Ile193Met variant at BTA1:94,396,804 bp (rs440830663) and **c** on the variant upstream *SPATA16* at BTA1:93,972,058 bp (rs379712951). The variants BTA1:94,396,804 and BTA1:93,972,058 are displayed in yellow and blue.

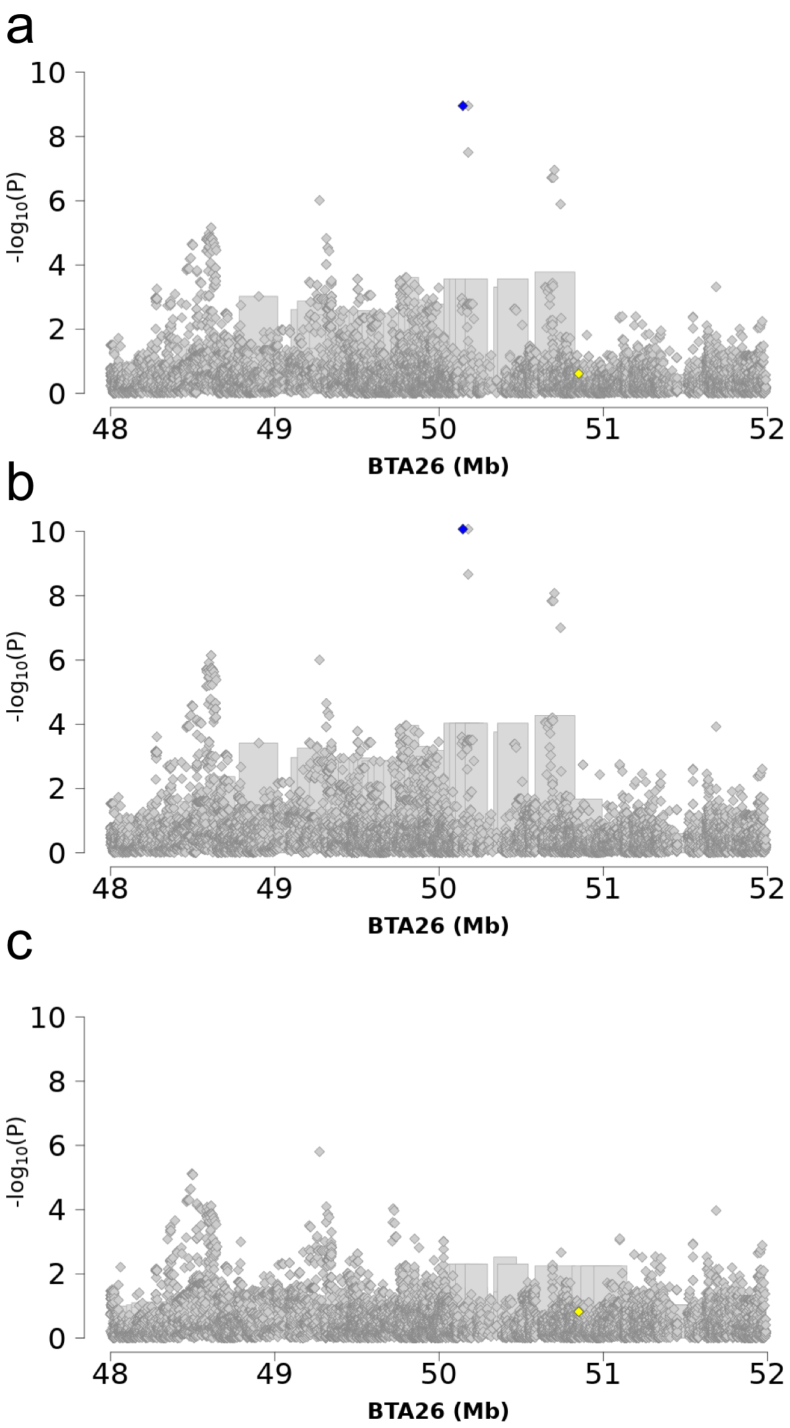


**Supplementary Figure 5: Conditional association analyses on BTA26 48-52 Mb. a** The imputed sequence variants (diamonds) and haplotypes (bars) association analyses were conditioned on the top haplotype of the BTA26 QTL, **b** on the missense variant BTA26:50,850,915C>G (rs378141069) in *ENSBTAG00000019919*, and **c** on the BTA26:50145932C>T (rs720936782) variant. The variants BTA26:50,850,915C>G and BTA26:50145932C>T are displayed in yellow and blue.

### Supplementary File

**Supplementary File 1: Accession numbers of 125 BSW animals.** The numbers listed indicate accession numbers from the sequence read archive of the European Nucleotide Archive (http://www.ebi.ac.uk/ena). The status of the top haplotypes for the QTL on BTA1, 18, 25, and 26 are given as 0, 1, and 2 (non-carrier, heterozygous, and homozygous). Bull fertility is given in the last column.

### Supplementary Data

**Supplementary Data:** Results of haplotype-based association testing for bull fertility, six semen quality, and 22 sperm morphology features. Results of sequence-based association studies for bull fertility at the BTA1, BTA18, BTA25 and BTA26 QTL.
